# Neuronal activity promotes repair by microglial reprogramming following demyelination

**DOI:** 10.1101/2025.11.19.689194

**Authors:** C Perrot, R Ronzano, Z Li, MS Aigrot, V Pantazou, P Stheneur, FX Lejeune, B Zalc, F Quintana, B Stankoff, C Lubetzki, A Desmazieres

## Abstract

Microglia, the resident immune cells of the central nervous system (CNS), play a multifaceted role in neurological disorders. In multiple sclerosis (MS), a chronic demyelinating and neurodegenerative disease, microglia contribute to inflammation and tissue damage, but can also support repair by clearing myelin debris, limiting inflammation and promoting remyelination and neuroprotection. The timely transition from their pro-inflammatory to pro-regenerative states is essential for effective repair and, in chronic MS, persistent, defective microglial activation contributes to disease progression. Yet, the mechanisms underlying the microglial switch remain largely unknown. In this study, we demonstrate that neuronal activity can modulate microglial signature at the onset of remyelination in MS models, in a pattern-dependent manner. Transcriptomic analyses reveal a downregulation of pro-inflammatory, disease-associated microglial signatures alongside an upregulation of genes associated with oxidative phosphorylation and lipid metabolism, indicative of a shift toward pro-regenerative states following physiological activity enhancement. This activity-dependent reprogramming also extends to infiltrating monocytes and macrophages, collectively fostering a microenvironment favoring repair.

## Introduction

Microglia are the main resident immune cells of the central nervous system (CNS), where they play crucial roles in neurodevelopment, homeostasis, and response to infection or injury ^1–4^. In multiple sclerosis (MS), a chronic inflammatory, demyelinating and neurodegenerative disease of the CNS, microglia can exhibit a dual role. They can either exacerbate pathology by promoting tissue damage or support neuroprotection and remyelination. Recent transcriptomic studies have highlighted the heterogeneity of activated microglia in MS, revealing a spectrum of signatures ranging from pro-inflammatory to pro-regenerative ^5–9^, which reflects lesional and microenvironment variability in MS ^10–12^. As early responders, microglia rapidly undergo morphological and phenotypic changes, initially amplifying inflammation. Over time, however, they can contribute to tissue repair by clearing myelin debris, restraining inflammation, supporting neuronal survival, and promoting oligodendrocyte differentiation and remyelination ^4,13,14^. This pro-regenerative state is characterized by transcriptomic reprogramming and a metabolic shift from glycolysis to oxidative phosphorylation. In MS, however, this transition can be impaired, leading to persistent pro-inflammatory states. This dysfunction contributes to chronic inflammation, oxidative stress, iron accumulation, ongoing demyelination and axonal damage, thereby driving disease progression ^8,15–17^.

In line with their specialization in CNS surveillance, microglia continuously integrate a wide range of signals, which modulate their behavior. In MS and its models, lymphocytes, in particular Th1 and Th17 T cells, can promote pro-inflammatory microglial phenotypes through the secretion of factors such as IFNγ, IL-17 and TNF- α leading to microglial cytokine release and antigen presentation ^18–20^. In contrast, pro-regenerative signals such as IL-4, IL-10 and TGF-β, secreted by Th2 cells and Tregs can orientate microglia towards reparative states ^21,22^.

Furthermore, microglia are sensitive to neuronal secreted factors, such as BDNF, which plays an important role in crosstalk modulating neuroinflammation and neuroprotection ^23,24^. Microglia also sense neuronal activity through detection of activity-released factors such as potassium, ATP or neurotransmitters ^25–30^. Neuronal activity further plays a crucial role in regulating myelination and is also an established modulator of remyelination, playing a direct role on OPC recruitment, proliferation, differentiation and myelin deposition ^31–37^. Whether it may also modulate microglia to promote repair following demyelination has not been shown so far.

Here, we demonstrate that neuronal activity plays a key role in reprogramming microglial phenotypes at the onset of remyelination. Through chemogenetic and optogenetic approaches applied in *ex vivo* and *in vivo* mouse models, we show that enhancing neuronal activity at the onset of repair promotes neuron-microglia communication and modulates microglial profiles, in a pattern-dependent manner. Using transcriptomic analyses, including single-cell RNA sequencing, we further show that stimulating neuronal activity *in vivo* during the early phase of repair following a toxic-induced demyelination induces significant microglial transcriptomic changes. These include a downregulation of genes related to disease-associated microglia (DAM) signatures and an upregulation of genes associated with oxidative phosphorylation, indicative of a switch towards pro-regenerative microglial states. Additionally, we observe a modulation of perilesional homeostatic microglia signature, with an upregulation of *Igf-1* and genes involved in lipid pathways. These pro-repair changes further extend to monocytes and macrophages, highlighting the strong influence of neuronal activity in creating a microenvironment permissive to remyelination, therefore opening novel therapeutic perspectives.

## Results

### Neuronal activity modulates microglial signature at the onset of repair in a pattern-specific manner ex vivo

The regulation of the oligodendroglial lineage by neuronal activity in repair is well documented, but how it impacts microglial function in remyelination is not known. To address this question, we first used a chemogenetic DREADDs approach to modulate neuronal activity in organotypic cultures of cerebellar slices, the virally induced expression of receptor hM3D(Gq)-mCherry leading to increased Purkinje cell activity upon clozapine-N-oxide ligand (CNO) addition to the culture medium (Fig. S1a).

We first assessed whether enhancing activity reinforces neuron-microglia interaction at the node of Ranvier, their preferential site of contact along axons ^29^. A 1-hour enhancement of Purkinje cell activity using DREADDS in myelinated slices led to a significant increase in neuron-microglia contact at nodes of Ranvier compared to vehicle-treated control slices, without altering nodal and microglial density or mean microglial total process length (Fig. S1b). We similarly observed an increase in microglia-node interaction after neuronal activity reinforcement at the onset of remyelination (2 days after LPC treatment, Fig. 1a), following lysolecithin (LPC) induced demyelination in cultured cerebellar slices.

**Fig. 1.**
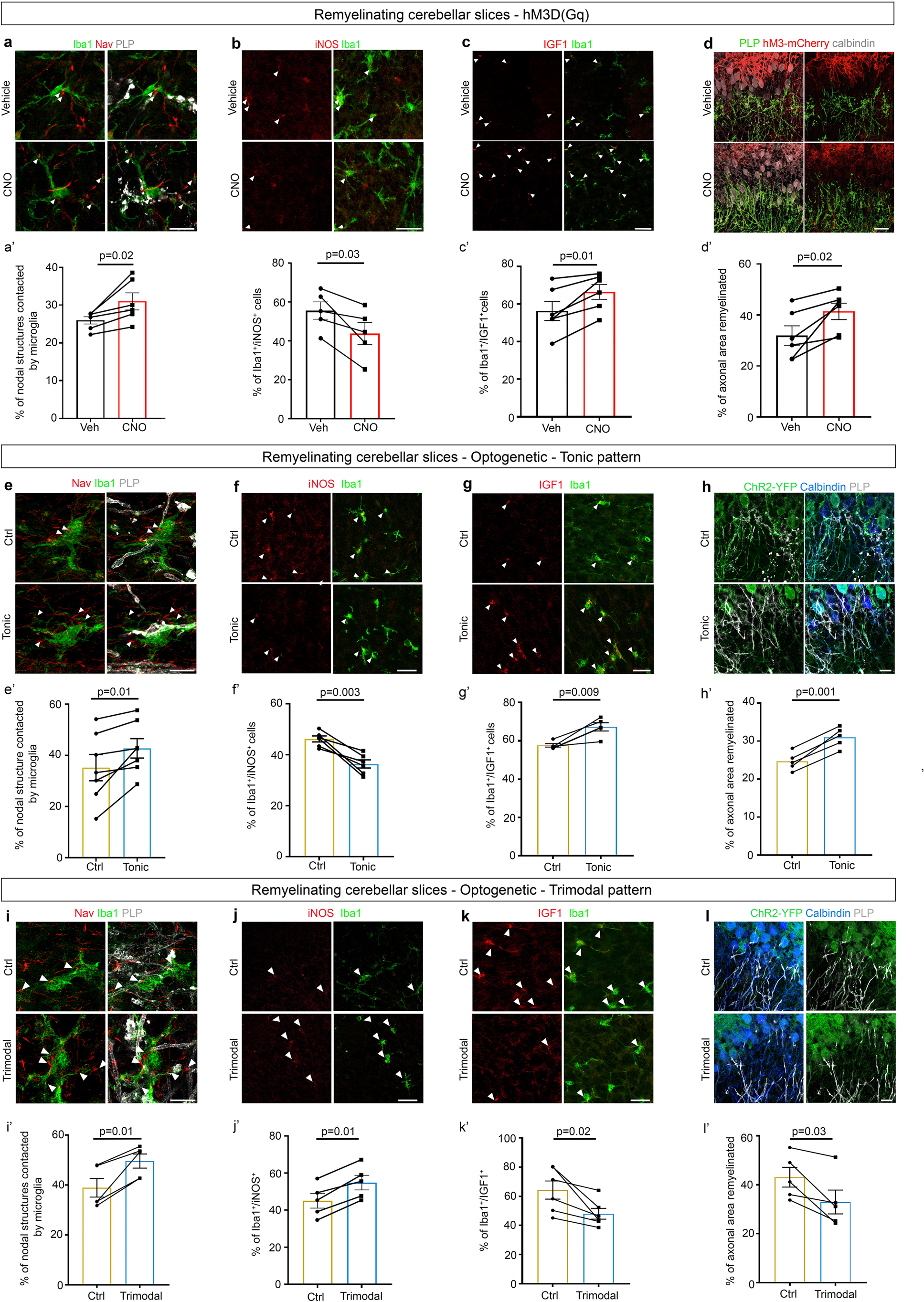
Neuronal activity modulates microglial phenotype and remyelination in a pattern-specific manner *ex vivo*. ***a–d****′,* Mouse cerebellar slices transduced with AAV8-hSyn-hM3D(Gq)-mCherry were demyelinated (LPC, 6-7 DIV) before being treated at the onset of remyelination (10 DIV) with CNO or its vehicle (DMSO) and fixed after 1 h (***a***) or 6 h **(*b–c*).** To quantify remyelination rate, the slices were treated for 6h and fixed 10h after the end of the treatment (***d***). ***a***, Microglia (Iba1, green) contact nodal structures (Nav, red) in remyelinating areas (PLP, grey); arrowheads indicate microglia–node contacts. ***a****′,* Percentage of nodes contacted by microglia. ***b–c***, Expression of iNOS **(*b***, red) or IGF1 (***c***, red) in microglia (Iba1, green) is indicated by white arrowheads. ***b′–c′,*** Percentage of microglial cells expressing iNOS or IGF1. ***d***, Remyelination (PLP, green) of Purkinje cell axons (Calbindin, blue) in transduced areas (mCherry, red). ***d′***, Quantification of axonal area covered by myelin. ***e–h′***, Slices from L7-ChR2-EYFP mice were demyelinated at DIV6 and stimulated at DIV10 with a tonic light pattern (470 nm, Tonic or 590 nm, Ctrl). Slices were fixed at 1 h (***e***), 6 h (***f–g***) or 10 hours following the end of a 6h treatment **(*h*). *e***, Microglia (Iba1, green) contact nodal structures (Nav, red) in remyelinating areas (PLP, grey); arrowheads indicate microglia–node contacts. ***e′***, Percentage of nodes contacted by microglia. ***f–g***, Expression of iNOS (***f*)** and IGF1 (***g***) in Iba1⁺ microglia (arrowheads). ***f′–g′***, Percentage of Iba1⁺ cells positive for iNOS or IGF1**. *h***, Remyelination (PLP, grey) of Purkinje cell axons (Calbindin, blue) expressing ChR2–YFP (YFP, green). ***h′***, Quantification of the remyelinated area**. *i–l′***, Slices were stimulated with a trimodal light pattern (470 nm, Trimodal or 590 nm, Ctrl) and fixed at 1 h (***i*)**, 6 h **(*j–k***) or 10h after the end of a 6h treatment **(*l*).** *i,* Microglia (Iba1, green) contact nodal structures (Nav, red) in remyelinating areas (PLP, grey); arrowheads indicate microglia–node contacts. ***i′***, Percentage of nodes contacted by microglia. ***j–k***, iNOS (***j***, red) and IGF1 (***k***, red) expression in Iba1⁺ microglia (arrowheads). ***j′–k′,*** Percentage of iNOS⁺ and IGF1⁺ microglia. ***l***, Remyelination (PLP, grey) of Purkinje cell axons (Calbindin, blue) expressing ChR2–YFP (YFP, green). ***l′***, Quantification of the remyelinated area. All data are presented as mean ± s.e.m. Statistical comparisons were performed using paired *t*-tests; full statistics are provided in Supplementary Table. Scale bars: ***a, e, i***, 10 µm**; *b–c, f–g, j–k*,** 20 µm**; *d, h, l*,** 30 µm.

As inhibiting microglia-neuron crosstalk at nodal structures following demyelination maintains pro-inflammatory microglia ^29^, we next asked whether we could reciprocally promote the switch towards pro-repair microglial phenotypes by increasing neuronal activity at the onset of remyelination. Slices in which neuronal activity had been stimulated for 5 hours at the onset of repair presented a 21% reduction in microglial cells expressing the pro-inflammatory marker iNOS, while the microglial pool expressing the pro-remyelinating factor IGF1 significantly increased by 1.2 folds compared to control slices (Fig. 1b–c). This shift in microglial marker expression correlated with increased remyelination, as measured by the axonal area (Calbindin, blue) covered by myelin (PLP^+^, grey), with a 30% increase in myelinated axonal area in slices with enhanced neuronal activity compared to controls (Fig. 1d).

We next assessed whether the firing pattern affects microglia-node contact and microglial signature in remyelination. To do so, we performed an optogenetic stimulation of Purkinje cell activity using L7-ChR2-YFP mice cerebellar slices, where channelrhodopsin is expressed specifically in Purkinje cells. The slices were stimulated using blue light (470 nm wavelength), a stimulation with yellow light (590nm wavelength) being used as control condition. To induce a physiological increase in Purkinje cell activity, we used a 10 Hz tonic stimulation pattern, while, for pathological stimulation, we used a trimodal illumination pattern leading to a baseline tonic firing, switching to high frequency firing followed by silent periods. This trimodal pattern mimics abnormal activity pattern previously observed in electrophysiological recordings of acute cerebellar slices from an inflammatory MS mouse model ^38^. Both paradigms induced a similar mean firing rate, as measured by patch-clamp recordings, thus differing only in their firing patterns (Fig. S1c–f). In myelinated as well as remyelinating cultured cerebellar slices, both stimulation profile led to a significant increase in the percentage of nodes contacted by microglia (Fig. 1e and i), without significantly altering microglial cell morphology, nodal density and microglial cell number (Fig. S1g–h and Fig. 1e-i).

They however differentially modulated microglial markers at the onset of repair. A tonic stimulation for 5 hours led to a significant 21% reduction in iNOS^+^/Iba1^+^ cells and a significant 17% increase in IGF1^+^/Iba1^+^ cells, associated to improved remyelination (26% increase in myelinated axonal area, Fig. 1f–h), confirming the results previously obtained with DREADDs-induced stimulation. In contrast, the trimodal pattern stimulation led to a 25% decrease in IGF1^+^ microglial cells and a 22% increase in iNOS^+^ microglia compared to control condition, associated to a 24% reduction in remyelination (Fig. 1j–l).

As high-frequency firing bursts can be associated to extracellular ATP release, which can be perceived by microglia ^39^, we hypothesized that the alteration of the microglial switch observed following trimodal pattern stimulation could be associated to purinergic signaling. P2X7 is a purinergic receptor highly expressed in activated microglia, where it is predominantly linked to pro-inflammatory functions. Pharmacological inhibition of P2X7 during trimodal stimulation of remyelinating cerebellar slices led to a significant rescue of the alteration in microglial phenotype induced by the trimodal stimulation (Fig. S1i–j), suggesting that purinergic signaling is implicated in aberrant pattern sensing by microglia.

Thus, neuronal activity modulates microglia behavior in repair by reinforcing microglia-node interaction and triggering phenotypic evolution. While a physiological enhancement of neuronal activity promotes the switch towards pro-regenerative microglial signatures and remyelination, aberrant activity maintains an inflammatory microglial polarization, identifying microglial cells as efficient sensors of neuronal activity dynamics in repair.

### Neuronal activity modulates microglial phenotype at the onset of remyelination in vivo

We next developed an *in vivo* approach to study the effect of neuronal activity on microglial signature at the onset of remyelination in an integrated context. Retrograde AAV vectors driving the neuronal expression of hM3D(Gq)-mCherry or hM4D(Gi)-mCherry were injected into the dorsal funiculus of mouse spinal cord, enabling neuronal activity activation or inhibition respectively in the dorsal funiculus of the spinal cord, while an AAV driving the expression of mCherry was used to assess the innocuity of CNO injection.

The spinal cord from mice transduced with AAVrt-hSyn-mCherry and treated with CNO presented no significant differences in the percentage of nodes contacted by microglia, node density and microglial number or process length compared to the mice transduced with AAVrt-hSyn-mCherry and treated with NaCl (Fig. S2a and S2b), showing CNO does not exert an effect on the studied mechanism per se. In myelinated spinal cord of hM3D(Gq)-expressing mice, a 1-hour CNO administration increased the proportion of nodes which are contacted by microglia by 33% without altering node density, microglial cell number, or average process length (Fig. S2c). Conversely, neuronal inhibition via hM4D(Gi) decreased microglial contacts at nodes by 25% (Fig. S2d). For the remyelinating condition, neuronal activity was modulated at the onset of repair, 2 days after the peak of LPC-induced focal demyelination, by systemic administration of clozapine-N-oxide (CNO) (Fig. 2a). Following a 1-hour modulation of neuronal activity, we observed a modulation of nodal contact similar to what we observed in the myelinated condition, while node density, microglia number and process length were not significantly changed between stimulated and control condition (Fig. 2b, hM3-Cherry and 2f, hM4-Cherry and Statistical table). These findings indicate that neuronal activity dynamically regulates microglial surveillance at nodal domains *in vivo*.

**Fig. 2.**
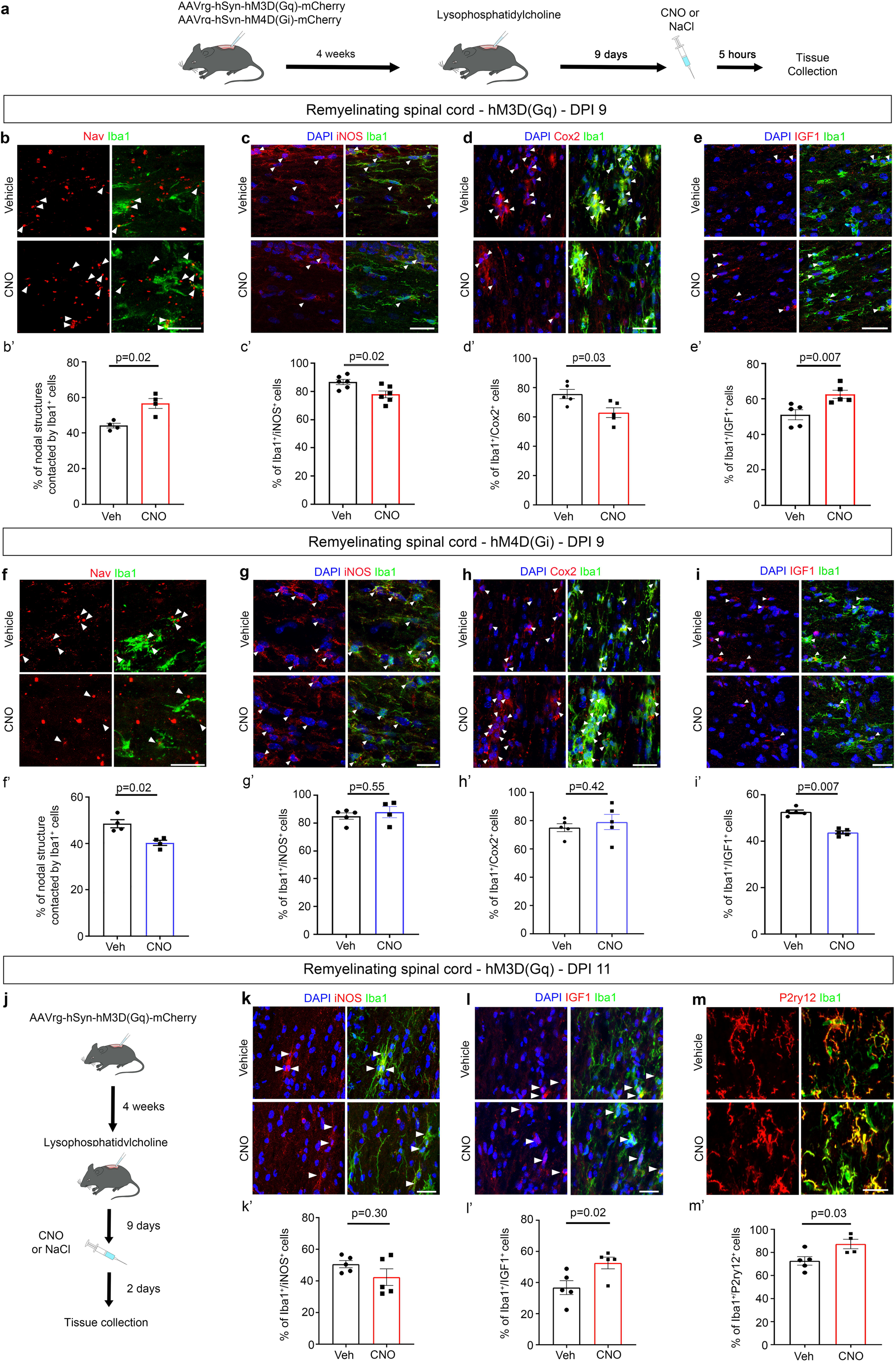
Neuronal activity modulates microglial phenotype at the onset of repair *in vivo*, with long-lasting effects. **a**, Experimental design for neuronal activity modulation in mouse spinal cord following demyelination. Tissues were collected 1 h (**b**) or 5 h (**c-e**, **j–m**) after neuronal activity enhancement (CNO) vs no activity modulation (Vehicle) at the onset of repair. **b,** Spinal cord sections immunolabeled for microglia (Iba1, green) and nodal structures (Nav, red); arrowheads indicate microglia–node contacts. **b′,** Percentage of nodal structures contacted by microglia. **c–e,** Iba1⁺ cells (green) co-expressing iNOS (**c**), Cox2 (**d**) or IGF1 (**e**) (red) are indicated by arrowheads; nuclei stained with DAPI (blue). **c′–e′,** Quantification of Iba1⁺ cells expressing iNOS, Cox2 or IGF1. **f–i,** Tissues were collected 1h (**f**) or 5 h post-treatment (**g-i**). **f,** Spinal cord sections immunolabeled for microglia (Iba1, green) and nodal structures (Nav, red); arrowheads indicate microglia–node contacts. **f′,** Percentage of nodal structures contacted by microglia. **g–i,** Iba1⁺ cells (green) co-expressing iNOS (**g**), Cox2 (**h**) or IGF1 (**i**) (red) are indicated by arrowheads; nuclei stained with DAPI (blue). **g′–i′,** Percentage of Iba1⁺ cells expressing iNOS (**g′**), Cox2 (**h′**), or IGF1 (**i’)**. **j–m′,** Spinal cord tissues were collected 2 days after stimulation (11 dpi) to assess the long-term effect of enhanced neuronal activity. **k–m,** Iba1⁺ cells (green) co-expressing iNOS (**k**), IGF1 (**l**) or P2RY12 (**m**) (red) are indicated by arrowheads; nuclei stained with DAPI (blue). **k′–m′,** Percentage of Iba1⁺ cells expressing iNOS, IGF1, or P2RY12. Data are presented as mean ± s.e.m. Comparisons were performed using Mann–Whitney tests (all panels), except **k** and **l** (unpaired *t*-tests). Scale bars: **b, g,** 10 µm; **c–e, h–i, k–l,** 20 µm.

A 5h-stimulation of neuronal activity at the onset of remyelination further led to a reinforced microglial phenotypic shift toward a pro-remyelinating state compared to control condition, with a significant reduction in Iba1^+^ cells expressing the pro-inflammatory markers iNOS and Cox2, while IGF1^+^/Iba1^+^ pool increased by 22% (Fig. 2c–e). This polarization was accompanied by enhanced microglial process complexity, as assessed by Sholl analysis and process length measurements (Fig. S3a). Neuronal activity inhibition, in contrast, led to a 17% reduction of IGF1^+^/Iba1^+^ population, while iNOS and Cox2 levels remained unchanged (Fig. 2g-i) and microglial morphology was only modestly affected (Fig. S3b).

### The microglial potassic channel THIK-1 is required for the modulation of microglia-node contact and microglial marker expression by neuronal activity at the onset of repair

The two-pore domain potassium channel THIK-1 was previously identified as a key player in microglia–neuron interaction at the node of Ranvier ^29,40^. We thus used THIK-1 KO mice ^41^ to assess whether it was required for the regulation of microglial signature by neuronal activity. We first confirmed that THIK-1 KO mice display a significant reduction in the proportion of microglia-node contact in the myelinated spinal cord, with no alteration in nodal and microglial density as well as myelination, but a mild reduction of microglial complexity (Fig. S4a-b). THIK-1 KO mice further showed a 35% reduction in nodes contacted by microglial processes at the onset of remyelination following LPC-induced demyelination (Fig. S4c), accompanied by a significant increase in CD68^+^ microglia (phagocytic marker, Fig. S4d), and reduced expression of iNOS^+^ and IGF1^+^ in THIK-1 KO microglia compared to control (Fig. S4e–f). In the absence of THIK-1, enhancing neuronal activity using DREADDs at the onset of remyelination failed to increase the frequency of microglia-node contacts and did not lead to a significant phenotypic evolution of microglial cells (Fig. S4g-j). These findings indicate that THIK-1 is required for microglial sensing of neuronal activity at nodes and for the associated microglial phenotypic modulation along spinal axonal tracts during remyelination.

### A transient neuronal activity stimulation at the onset of repair exerts durable effects on microglial phenotype and remyelination in vivo

To determine whether transiently enhancing neuronal activity at the onset of repair led to lasting changes in microglial state and in remyelination, we next assessed microglial signature 48 hours after the chemogenetic stimulation performed at 9 dpi (Fig. 2j). At this timepoint, neuronal activity impact was still visible, with significantly higher IGF1^+^ microglial rate (Fig. 2l) and a 20% increase of P2RY12^+^ microglia, a marker of homeostatic microglia, in the stimulated condition compared to controls (Fig. 2m), while iNOS+ microglia were mostly unaffected compared to control condition (Fig. 2k). This correlated with a significantly reduced lesion (PLP^-^/Iba1^+^ area, Fig. S3c) and a significant increase in node of Ranvier density in the lesional area (Fig. S3d), consistent with enhanced remyelination following a transient increase in neuronal activity at the onset or repair. These findings indicate that a transient increase of neuronal activity during the early phase of repair can durably shape microglial phenotype, correlating with enhanced remyelination.

### Transcriptomic analysis by RNA sequencing reveals a reprogramming of activated microglia by neuronal activity in repair

To further investigate how neuronal activity shapes microglial signature at the onset of repair, we performed transcriptomic analyses of mouse spinal cord with LPC-induced demyelinated lesion, following neuronal activity modulation at the onset of repair.

In an initial exploratory approach, we performed bulk RNA-sequencing and first confirmed that CNO administration does not trigger significant transcriptional changes, using AAVrt-hSyn-mCherry viral transduction coupled to CNO intraperitoneal injection. Indeed, we observed an overlap in the Principal Component Analysis (PCA) between mCherry/CNO and mCherry/NaCl samples (Fig. S5a) and no major gene expression modulation between the two conditions (Fig. S5d).

We next focused on the impact of neuronal modulation mediated by hM3D(Gq) or hM4D(Gi) activation. PCA revealed distinct segregation between stimulated and control conditions in both hM3D(Gq)-mCherry and hM4D(Gi)-mCherry expressing animals (Fig. S5b-c), indicative of robust transcriptomic differences between conditions. Increased neuronal activity resulted in the significant downregulation of 291 genes (Fig. S5e) and the significant upregulation of 19 genes. Pathway analysis using Ingenuity Pathway Analysis (IPA) revealed reduction in inflammatory cascades, including the downregulation of pathways such as the strongly pro-inflammatory “JAK/IL-6” signaling pathways, as well as “Macrophage activation” pathways (Fig. S5g). The expression of canonical inflammatory mediators such as *Ccl3* and *Cd9*, markers expressed in Disease Associated Microglia (DAM, ^42,43^) and peripheral immune cells, was significantly decreased (Fig. S5i–j). Furthermore, neuronal activity inhibition induced the upregulation of over 750 genes, while 120 were significantly downregulated (Fig. S5f). Upregulated genes included *Tnf*, *C1qa* and *Ccl5,* and reinforced inflammatory pathways such as “Neuroinflammation Signaling,” “Classical macrophage pathways” or “Complement activation” (Fig. S5h, k–l). All together, these findings support a beneficial role of neuronal activity in limiting tissular inflammation at the onset of repair.

In order to study transcriptomic signatures at the cellular level, we next performed single-cell RNA sequencing on mouse spinal cords following DREADD-induced neuronal activity modulation at the onset of repair (Fig. S6a). Quality control confirmed high-complexity libraries with adequate ribosomal RNA content and a median of 1000 genes per cell (Fig. S6b-d). Dimensionality reduction and unsupervised clustering identified 12 transcriptionally distinct cell populations, including microglia and monocytes/macrophages clusters (n = 92601 and n = 8238 cells, respectively; Fig. S6e-f; Supplementary Table 1).

Focusing on differentially expressed genes in the microglial cluster, we identified 313 genes significantly upregulated following activity stimulation (hM3D(Gq)/CNO vs hM3D(Gq)/NaCl; Fig. 3a-b). Following activity enhancement, microglia exhibited decreased expression of DAM genes such as *Clec7a*, *Hif1α*, *Lpl* or *Spp*1, and the inflammatory gene *Nfκb1*, indicating signatures with reduced inflammatory and oxidative stress profiles (Fig. 3c and Figure S7c). Some immune signaling pathways, including “NF-κB regulation” and “Cytokine-mediated responses” were downregulated as observed by Gene ontology enrichment analysis of the downregulated genes (GO Biological Process 2025; Fig. S7b; Supplementary Table 2), consistent with less inflammatory microglial signatures following neuronal activity enhancement.

**Fig. 3.**
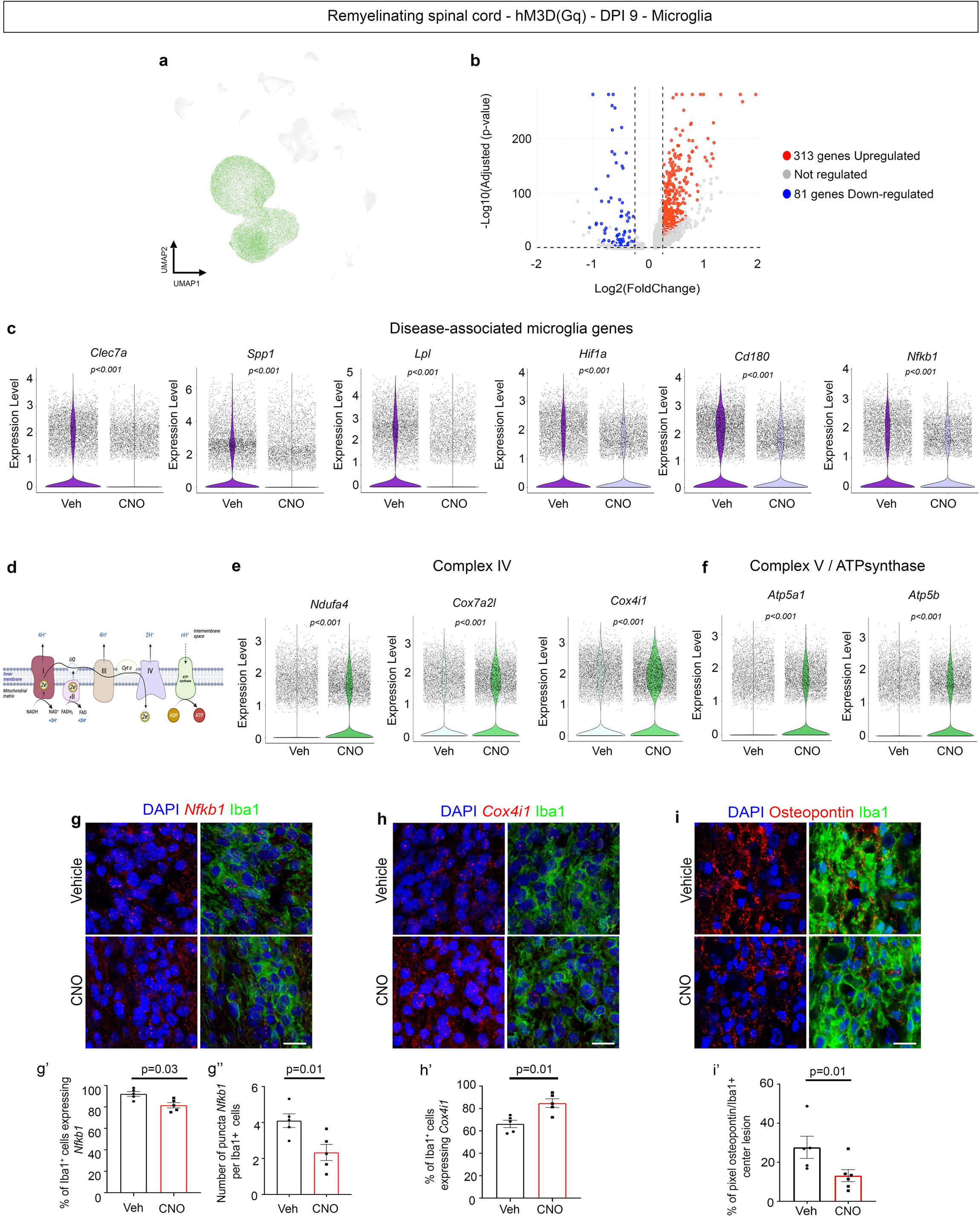
Neuronal activity promotes a transcriptomic shift in microglial cells at the onset of repair. **a**, UMAP representation of spinal cord cells isolated 5 h after CNO or vehicle administration in hM3D(Gq)-expressing mice at 9 days post-lesion (dpi). Microglia (green) were selected for downstream transcriptomic analyses. **b,** Volcano plot showing differentially expressed genes in microglia between CNO and vehicle (NaCl) conditions. A total of 313 genes were significantly upregulated and 81 genes downregulated (log₂ fold change > 0.25 or < –0.25; FDR < 0.05). **c,** Violin plots showing reduced expression of disease-associated and pro-inflammatory genes (*Clec7a, Spp1, Lpl, Hif1a, Cd180, Nfkb1*) in microglia from CNO-treated compared to control mice. **d,** Schematic representation of the mitochondrial electron transport chain. **e–f,** Violin plots showing expression levels of selected genes associated with respiratory chain complexes in microglia from CNO- or vehicle-treated mice. **g–i,** Representative spinal cord sections showing microglia (Iba1, green), DAPI (blue) and RNAscope signal (red) for **g,** *Nfkb1*, **h,** *Cox4i1*, and immunohistochemistry for Osteopontin, **i**. **g’-g’’,** percentage of Iba1⁺ cells expressing *Nfkb1* and number of *Nfkb1* puncta per Iba1⁺ cell. **h′,** percentage of Iba1**⁺**cells expressing *Cox4i1⁺*and **i′,** quantification of Osteopontin^+^ pixels within Iba1⁺ cells. Data are presented as mean ± s.e.m. Statistical comparisons were performed using Mann–Whitney tests. Scale bars: **g–i,** 10 µm.

Gene ontology enrichment analysis of the 313 upregulated genes further revealed a strong increase in oxidative phosphorylation (OXPHOS) and mitochondrial energy related pathways, including “Cellular respiration”, “Mitochondrial ATP synthesis”, and “Aerobic electron transport chain” (Fig. S7a). We further observed a significant upregulation of mitochondrial complex IV and V associated genes (such as *Cox4i1, Cox7a2i, Ndufa4* and *Atp5a1, Atp5b* respectively; Fig. 3d-f; Fig. S7c). These data suggest that neuronal activity promotes a shift towards aerobic OXPHOS metabolism in microglia (Fig. 3d-f; Fig. S7a-c), the switch from glycolytic to OXPHOS metabolism being classically associated to the transition from pro-inflammatory to pro-regenerative innate immune cells ^44–46^. In addition, genes involved in glutamate uptake (*Slc1a3*) and in the regulation of mitophagy and reactive oxygen species (*Fkbp5*, *Ucp2*) were robustly upregulated (Supplementary Table 2).

On the contrary, the inhibition of neuronal activity at the onset of repair led to the upregulation of 131 transcripts in microglia (hM4D(Gi)/CNO vs hM4D(Gi)/NaCl), with more than half corresponding to ribosomal genes, suggesting a deteriorated cellular health (71/131; Supplementary Table 3; Fig. S7d). After exclusion of these genes, enrichment analysis revealed activation of “Oxidative damage response” and “Classical complement cascade” pathways, with increased microglial expression of inflammation-associated genes such as *Cd74*, *Lyz2* or *H2-D1*. Concomitantly, transcripts associated to ATPase (*Atp2c1*) and TGF-β signaling (*Tgfbr2*) were downregulated (Fig. S7e–f).

To validate these results, we performed RNAscope analysis on spinal cord sections for specific targets. After 5 hours of increased neuronal activity, the proportion of microglia expressing *Cox4i1* was elevated by 27% in the lesion (Fig. 3h), while the proportion of *Nfκb1*-positive microglia was reduced by 12%, and the number of puncta per cell decreased by 78% (Fig. 3g). A 2.1-fold decrease in Osteopontin protein (encoded by *Spp1*) levels in Iba1^+^ cells following neuronal activity stimulation was further observed within the lesion (Fig. 3i), whereas neuronal activity inhibition enhanced its expression (53% increase, Fig. S7h).

Altogether, these findings indicate that neuronal activity drives microglia toward a less pro-inflammatory transcriptomic profile, characterized by reduction of DAM-related programs and enhanced mitochondrial oxidative phosphorylation. In contrast, neuronal activity inhibition leads to a pro-inflammatory, stress-associated profile.

### Neuronal activity stimulation leads to less pro-inflammatory and more pro-regenerative signatures in *activated microglial subclusters*

To further resolve whether neuronal activity modulates microglia globally or rather acts on specific microglial subpopulations during remyelination, we subclustered the microglial cells of the single-cell RNA sequencing dataset (Fig. 4a). Unsupervised clustering identified six transcriptionally distinct clusters, with a homeostatic microglial population (Cluster 1/HOM) and activated profiles including two subclusters with DAM-like signatures (Clusters 3/DL1 and 4/DL2, Cluster 3 showing a less pro-inflammatory signature than Cluster 4), one cluster transitioning to OXPHOS profile (Cluster 5/REG) and one cluster with a virally-induced profile (Cluster 6/VIR). Clusters 4 and 6, each comprising fewer than 5% of total microglia, were excluded from downstream analyses due to limited statistical power. Cluster 2 (SPL) was enriched for transcripts related to RNA processing (*Son, Ddx17*) and mitochondrial genes (*mt-Co1, mt-Nd1*) ^47^ and exhibited high expression of *Tet2* and *Tet3*, consistent with a metabolically and transcriptionally stressed signature (Fig. 4b–c; Supplementary Table 4) ^48^. In contrast to Clusters 3 (DL1), which expressed DAM-related genes, including *Clec7a*, *Spp1* and *Lpl*, and glycolysis-associated genes, such as *Gapdh* and *Mif*, Cluster 5 (REG) showed reduced expression of pro-inflammatory genes, including *Lpl*, *Spp1*, and *Hif1a* and a high expression of transcripts associated to oxidative phosphorylation (*Cox4i1, Cox8a*) and antioxidant defense (*Gpx1*, *Gpx4*), suggesting a transitioning, regenerative phenotype (Fig. 4b-c; Supplementary Table 4). Cluster 1 (HOM) presented a homeostatic microglia identity, marked by elevated *P2ry12* and *Tmem119* expression (Fig. 4b-c; Supplementary Table 4).

**Fig. 4.**
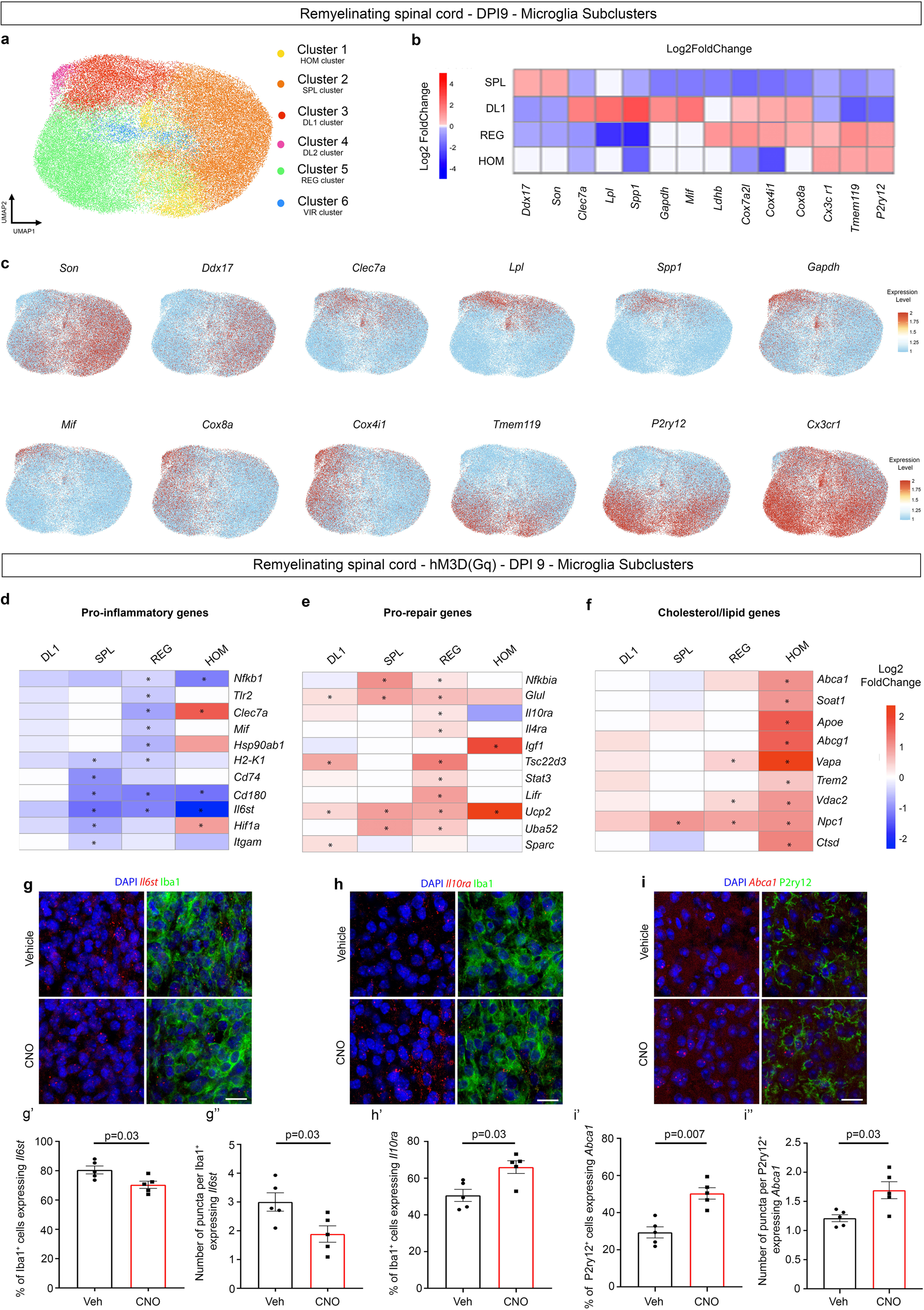
Neuronal activity induces a decrease in pro-inflammatory gene expression in activated microglia and an increase of lipid-related gene expression in homeostatic microglia. **a**, UMAP representation of microglial cells from the spinal cord of hM3D(Gq)- or hM4D(Gi)-expressing mice at 9 days post-lesion (dpi), 5 h after CNO or vehicle (NaCl) injection. Unsupervised clustering identified six transcriptionally distinct microglial clusters. **b,** Heatmap displaying relative expression (log₂ fold change) of selected marker genes across the four main clusters. Cluster 2 (SPL) signature was associated to RNA processing. Cluster 3 (DL1) exhibited a DAM-like signature (*Clec7a, Lpl, Spp1*), while cluster 5 (REG) showed features of a transitional pro-regenerative phenotype, with increased expression of *Cox4i1* and *Cx3cr1* and reduced DAM-associated transcripts. Cluster 1 (HOM) was enriched for homeostatic genes (*P2ry12, Tmem119*). **c,** UMAP feature plots showing expression of representative genes across clusters, including RNA-processing associated genes (*Son, Ddx17*), DAM-associated genes (*Clec7a, Spp1, Lpl*), glycolysis-associated genes (*Mif, Gapdh*), mitochondrial genes (*Cox4i1, Cox8a*) and homeostatic markers (*Tmem119, Cx3cr1, P2ry12*). **d–f,** Heatmaps showing differentially expressed genes between CNO- and vehicle-treated hM3D(Gq) mice across microglial clusters: **d,** pro-inflammatory genes, **e,** pro-repair genes, and **f,** cholesterol/lipid pathways-related genes. **g–i,** RNAscope combined with immunofluorescence in spinal cord lesion. **g,** *Il6st* RNAscope signal (red) in Iba1⁺ microglia (green). **g′–g″,** Percentage of *Il6st⁺* microglia and number of Il6st puncta per cell. **h,** *Il10ra* RNAscope (red) in Iba1⁺ cells (green). **h′,** Percentage of *Il10ra⁺* microglia. **i,** Confocal images showing *Abca1* expression (red) in P2RY12⁺ microglia (green). **i′–i″,** Percentage of P2RY12⁺ microglia expressing *Abca1* and number of puncta per cell. Data are presented as mean ± s.e.m. Mann–Whitney tests. Scale bars: **g–i,** 10 µm.

To capture the dynamic evolution of these microglial states, we applied pseudotime trajectory analysis using Slingshot (Figure S8) on DL1, REG and HOM subclusters, using the DAM-like cluster DL1 as the root. This revealed a continuous transcriptional trajectory progressing from disease-associated microglia to a pro-regenerative state, ultimately leading to the homeostatic profile (Fig. S8a). Along this trajectory, we observed a gradual downregulation of canonical DAM genes (*Lpl*, *Spp1*, *Clec7a*) and a concomitant upregulation of homeostatic markers including *Tmem119*, *Cx3cr1*, *P2ry12* and *Csf1r* (Fig. S8b).

To examine how neuronal activity modulates this progression, we compared transcriptional responses between stimulated and control conditions (hM3D(Gq)/CNO and hM3D(Gq)/NaCl) across clusters and along pseudotime. Analysis of the distribution of cells along the pseudotime trajectory revealed a higher proportion of cells at low-pseudotimes (<30), corresponding predominantly to DAM-like state, in the hM3D(Gq)/NaCl control condition (Fig. S8a-c). As pseudotime increased, we observed a shift in distribution, with a higher proportion of cells derived from the increased activity condition (hM3(Gq)/CNO). This transition occurred around pseudotime 35, a region characterized by transitioning, regenerative microglial states (Fig. S8c).

In cluster-based analysis, neuronal activation led to the downregulation of pro-inflammatory genes, such as *Nfκb1*, *H2-K1* or *Il6st* (Fig. 4d; Supplementary Table 5) in activated subclusters, with a more potent effect on the REG cluster. This suppression was accompanied by a significant upregulation of transcripts involved in immune regulation (*Nfκbia, Il4ra, Il10ra*), glutamin/glutamate metabolism (*Glul*), ROS detoxification (*Ucp2*), and mitophagy (*Fkbp5*) in all the activated subclusters (Fig. 4e; Supplementary Table 5), all consistent with the emergence of metabolically competent and immuno-regulatory microglial states. Furthermore, the REG cluster specifically presented a significant upregulation of *Lifr*, which is associated to inflammation regulation and neuroprotection ^49^. Using RNAscope, we observed a reduction of *Il6st* expression in the center of the lesion following neuronal activity stimulation, with a small reduction of *Il6st* expressing microglia, but a 37% drop in the number of puncta per microglial cell (Fig. 4g). In contrast, *Il10ra* expressing microglia were increased by 31% in this same area following neuronal activity stimulation (Fig. 4h). Differential pseudotime transcripts analysis between stimulated and control animals further revealed 69 genes significantly upregulated and 45 downregulated upon neuronal stimulation (Fig. S8d; Supplementary Table 6).

This study confirmed the positive effect of neuronal activity for the switch between pro-inflammatory and pro-regenerative microglial profiles, with DAM-associated genes such as *Spp1* and *Hif1α* expression being reduced in DAM-like states following neuronal activity stimulation (Fig. S8e-f), while neuroprotective and anti-inflammatory transcripts such as *Il10ra* were significantly increased in the later stages (Fig. S8g; Supplementary Table 6). *Fkbp5* and *Glul* were furthermore among the most enriched transcripts following activity enhancement in this analysis (Fig. S8i; Supplementary Table 6). Interestingly, we observed a marked upregulation of *Ddit4* expression in the stimulated condition at the pseudotime point corresponding to the microglial shift (Fig. S8j). *Ddit4* has previously been identified as a key regulator of macrophage polarization through its modulation of the mTOR pathway and of glycolytic metabolism ^50^. These findings suggest that *Ddit4* may act as a molecular switch for microglial reprogramming in response to neuronal activity.

### Neuronal activity orientates homeostatic microglia towards remyelination-promoting signatures

Given the major role of lipids in remyelination ^51–53^, we next assessed the expression of genes involved in lipid metabolism and transport in the various microglial subclusters. Surprisingly, homeostatic microglia showed a strong upregulation of lipid-associated genes in response to neuronal activity (Fig. 4f), including *ApoE* and *Trem2*, implicated in myelin debris scavenging, as well as *Abca1* and *Abcg1*, which mediate lipid efflux. Pseudotime analysis confirmed a mild but significant upregulation of *Abca1* in microglia occupying higher pseudotime values (>40), corresponding to a homeostatic phenotype, following activity stimulation (Fig. S8h). These findings were validated by a RNAscope study, showing a 72% increase in the proportion of P2RY12⁺ microglia expressing *Abca1* in the perilesional area and a 45% increase in puncta density per positive cell after neuronal activation (Fig. 4i). Reciprocally, neuronal activity inhibition led to downregulation of lipid-associated genes in homeostatic microglia (HOM, Fig. S9c), while it did not induce major changes in activated microglia (Fig. S9a-c). Following neuronal activity inhibition, a 1.93-fold reduction in the proportion of P2RY12⁺ microglia expressing *Abca1* was further observed by RNAscope (Fig. S9d).

Altogether, our results demonstrate that neuronal activity participate in microglial identity reprograming at the onset of repair, by downregulating inflammatory and DAM associated genes, attenuating mitochondrial stress, while enhancing OXPHOS metabolism, mitophagy and pro-regenerative gene expression, and regulating lipid metabolism and fluxes processes.

### Neuronal activity modulates inflammatory gene expression in monocyte and macrophage populations

To determine whether the effects of neuronal activity on the immune landscape extended beyond microglia, we next examined the impact of activity modulation on the monocyte/macrophage cluster identified in our scRNA seq study, which we could subdivide in five distinct subclusters (Fig. 5a). Clusters 1 and 5 were characterized by high *Ly6c2* and *F10* expression, consistent with monocytic profile, with cluster 5 further exhibiting elevated *Top2a* and *Mki67* levels, indicating a proliferative phenotype (Fig. 5b-c, Supplementary Table 7). In contrast, Cluster 2 displayed gene expression patterns indicative of a macrophage identity, including robust expression of *Ms4a7* and *Mrc1* (Fig. 5b-c, Supplementary Table 7). Clusters 3 and 4 were associated to dendritic cell and leukocytic profiles respectively (Supplementary Table 7).

**Fig. 5.**
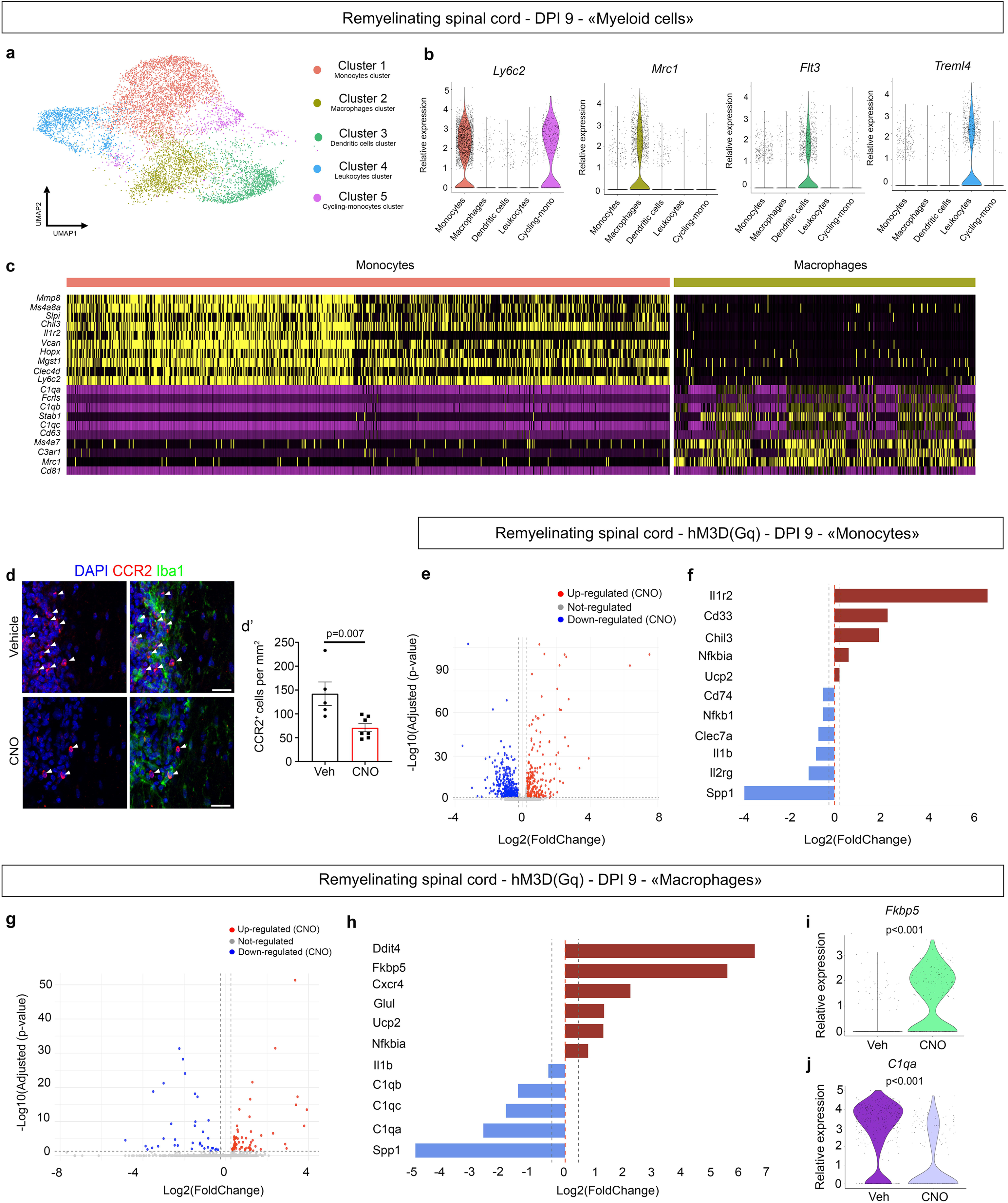
Neuronal activity reprograms monocyte and macrophages by reducing inflammatory signatures and promoting repair-associated gene expression. **a**, UMAP representation of myeloid cells in spinal cords of hM3D(Gq)-expressing mice at 9 days post-lesion (dpi), 5 h after CNO or vehicle treatment. Subclustering identified five transcriptionally distinct populations: cluster 0 (monocytes), cluster 1 (macrophages), cluster 2 (dendritic cells), cluster 3 (leukocytes), and cluster 4 (cycling monocytes). **b,** Violin plots showing expression of representative markers for monocytes (*Ly6c2*), macrophages (*Mrc1*), and other immune subsets. **c,** Heatmap displaying the top 10 differentially expressed genes for monocyte (left) and macrophage (right) clusters. **d,** Representative spinal cord section immunostained for Iba1⁺ (green) and CCR2⁺ (red) cells in the lesion. **d′,** CCR2⁺ Iba1⁺ cells were significantly reduced in CNO-treated mice compared to control. Data are presented as mean ± s.e.m. **e,** Volcano plot showing differentially expressed genes in the monocyte cluster between CNO- and vehicle-treated mice. A total of 198 genes were significantly upregulated and 317 downregulated (log₂ fold change > 0.25 or < –0.25; FDR < 0.05). **f,** Selected genes differentially expressed in monocytes. CNO treatment increased anti-inflammatory (*Il1r2, Cd33*) and stress-response (*Ucp2*) genes, and reduced expression of pro-inflammatory (*Il1b, Nfkb1*) and DAM-associated (*Clec7a, Spp1*) genes. **g,** Volcano plot of differentially expressed genes in the macrophage cluster between CNO and vehicle conditions. A total of 68 genes were significantly upregulated and 36 downregulated (log₂ fold change > 0.25 or < –0.25; FDR < 0.05). **h,** Among the genes modulated in macrophages, upregulated genes included *Fkbp5, Ddit4, Ucp2*, while downregulated genes included complement-associated genes (*C1qa, C1qb, C1qc*) and the DAM marker *Spp1*. **i–j,** Violin plots showing increased expression of *Fkbp5* (**i**) and reduced expression of *C1qa* (**j**) in macrophages from CNO-treated mice compared to controls. Mann–Whitney tests (**f′**). Scale bar: **f,** 20 µm.

We first observed that neuronal activity stimulation led to a significant 50% reduction in the number of peripheral innate immune cells following neuronal activity stimulation (CCR2^+^/Iba1^+^ cells, Fig. 5d). To assess how neuronal activity influences monocyte and macrophage transcriptomic signature, we performed differential expression analysis in stimulated vs control conditions for Cluster 1 and Cluster 2 respectively. In the “monocyte” cluster, 168 genes were significantly upregulated and 317 genes were downregulated in response to neuronal activity enhancement (Fig. 5e, Supplementary Table 8). The most strongly induced transcript was *Il1r2* (Fig. 5f), which has recently been identified as a negative regulator of monocyte recruitment during inflammation ^54^. Among the upregulated genes, we observed an enrichment in regulators of inflammatory pathways, including *Cd33*, a gene involved in the negative regulation of cytokine production. We also observed a downregulation of DAM-associated genes and inflammatory genes such as *Spp1*, *Clec7a* and *Nfκb1* and an increased expression in *Fakbp5* gene and *Glul*, as observed for microglia, as well as in transcripts associated to oligodendrogenesis support, such as *Chil3* (^55^; Supplementary Table 8).

In macrophages, neuronal activity stimulation led to a significant upregulation of 68 genes and a downregulation of 36 genes (Fig. 5g, Supplementary Table 8). We again observed a reduction of DAM-associated genes such as *Spp1*, inflammatory genes such as *Il1β*, as well as classical complement pathway components, including *C1qa*, *C1qb*, and *C1qc* (Fig.5h-j, Supplementary Table 8), which are implicated in particular in chronic neuroinflammation in MS ^8,56^. We also observed an increase in *Fkbp5*, *Ucp2*, *Glul* and *Ddit4* as seen for microglia (Fig. 5i-j, Supplementary Table 8).

These changes suggest that monocytes and macrophages can be reprogrammed by neuronal activity similarly to microglia, with neuronal activity enhancement leading in particular to increased expression of genes supporting the switch towards pro-repair phenotypes in macrophages.

## Discussion

In this work, we show that neuronal activity modulates microglial phenotypes at the onset of repair following demyelination, in a pattern-dependent manner. We further demonstrate that a physiological enhancement of neuronal activity promotes the switch towards regenerative microglia, with a similar effect on monocytes and macrophages.

Neuronal activity is a well-established regulator of the oligodendroglial lineage and myelin deposition in myelination, plasticity and repair ^32,33,57^. However, its influence on microglial behavior in remyelination is not clearly described. In MS and its models, it has previously been shown that physical exercise, which can enhance neuronal activity amongst other processes, can modulate remyelination and immunity ^58,59^. In MS models, though, these trainings were preventive or performed just after demyelination induction ^58,59^. Here, we asked whether enhancing neuronal after the peak of demyelination can modulate microglial phenotype and promote its switch towards a pro-regenerative status. We show that neuronal firing indeed serves as an instructive factor that modulates microglial phenotypic states, indirectly influencing the efficiency of remyelination. THIK-1, a major microglial potassium channel participating in neuron-microglia crosstalk at nodes ^29^, is furthermore essential for neuronal activity to effectively modulate microglial signature at the onset of repair.

We further assessed whether neuronal activity pattern could affect the modulatory effect of neuron-microglia communication on microglial signature. This question is of particular interest as abnormal firing patterns, including hyperactivity, have been observed in demyelinating models ^38,60^. We show that, while a physiological firing enhancement is beneficial regarding microglial signatures and increases remyelination, an activity pattern with high frequency bursts reinforces the pro-inflammatory state of microglia and impedes repair. This is consistent with previous studies showing that aberrant activity following demyelination is unfavorable for oligodendrocyte differentiation and remyelination ^61^. We next showed that purinergic signaling participates in abnormal pattern detection. Neuronal hyperactivity is known to promote the extracellular release of ATP, which can lead to the activation of microglial purinergic receptors ^39^. At high concentrations (∼100 µM), ATP activates P2X7, a receptor upregulated upon microglial activation and known to trigger NLRP3 inflammasome activity and pro-inflammatory gene expression ^62^. In contrast, activation of P2X4, a purinergic receptor sensitive to lower ATP concentration, has been linked to reparative functions ^63^, suggesting that ATP may act as a context-dependent signal whose effect depends on local concentration and receptor availability. Astrocytes, which can also contact nodes of Ranvier, could further contribute to this signaling cascade by releasing ATP in response to neuronal firing ^64^. Neuronal activity can further modulate the secretion of multiple neuronal cues, such as neurotransmitters as well as growth and immunomodulatory factors, including IL-13, which can participate in microglial activation and phenotypic orientation ^24,30,65^, thus suggesting the existence of a complex neuronal activity-dependent signaling shaping an adapted microglial behavior in response to neuronal physiological status.

We next demonstrated that neuronal activity can trigger a transcriptomic reprogramming of microglia at the onset of repair. Upon activity stimulation, genes associated to antigen presentation, glycolytic or inflammatory states (such as *Hk2-K1*, *CD180, Mif, Gapdh, Nfκb1, Spp1 or Hif1a,*) were downregulated, while genes associated to anti-inflammatory and regenerative pathways, such as *Il4ra*, *Il10ra* or *SPARC/Uba52* were upregulated. In parallel, we observed a significant upregulation of genes associated to oxidative phosphorylation (*Cox4i1, Nduf4a*), mitochondrial quality control (*Fkbp5, Ucp2*) and antioxydant response (*Gpx1* and *Gpx4*) in all the activated microglial subclusters studied (DL-1, SPL and REG). Gpx4 is further known to be a potent inhibitor of ferroptosis, a mechanism induced by iron accumulation which is at play in MS ^66^ and *Fkbp5* has been shown to be essential for establishing a pro-remyelinating environment ^67^. These changes may in particular restore redox homeostatis and limit ROS production by limiting reverse electron transport, a major source of oxidative damage in chronic inflammation ^68,69^. The upregulation of genes such as *Glul* and *Ldhb* may further support mitochondrial metabolism ^70,71^. Altogether, these data points to an engagement towards healthier, pro-regenerative microglial states. This is consistent with the previously observe shift from OXPHOS to glycolytic metabolism upon microglial activation, whereas pro-regenerative states depend on switching back towards oxydative respiration ^45,46,68,72^.

Furthermore, recent studies have implicated microglia in complex lipid handling during remyelination, contributing not only to debris clearance but also to the supply of cholesterol and lipids required for myelin synthesis ^52,73^. Here, we show that neuronal activity induces in homeostatic microglia a coordinated increase in the expression of *ApoE, Trem2, Abca1* and *Abcg1*, key genes driving lipid sensing, trafficking, and efflux. This may participate in the preservation of lysosomal integrity and prevent lipid overload, maintaining microglia in a competent, protective state. Indeed, it was recently shown that loss of lipid efflux genes lead microglia to deleterious foamy *Gpnmb*^+^ phenotype, associated with severe disease course ^8,17,74,75^. Such regulation suggests that neuronal signals actively sustain microglial lipid recycling, thereby maintaining cholesterol availability required for oligodendrocyte differentiation and myelin synthesis, a process often limited by cholesterol scarcity during repair ^51,52,73^. These findings bring out neuronal firing as a regulator of microglial lipid metabolism, suggesting neuronal activity could promote remyelination through a coordinated neuron–microglia–oligodendrocyte axis. The nature of the activity-induced pathways leading to microglial reprogramming of lipid associated genes remains to be described.

While microglia are the principal immune cells of the CNS, peripheral innate immune cells infiltrate demyelinated lesions, where they are rather considered to exacerbate damage ^76^. Our data show that neuronal activity also reprograms monocytes and macrophages within demyelinated lesions and induced a shift toward a reparative state, suppressing transcripts linked to complement signaling and neuroinflammatory activation (*C1qa, Spp1, Clec7a, Il1b*) ^8^, while genes promoting oligodendrogenesis and myelin integrity, including *Chil3* and *Tgfbr1* were upregulated ^55,77^. Interestingly, the expression of *Il1r2*, which encodes a potent inhibitor of monocyte recruitment during inflammation ^54^, was strongly upregulated, consistent with the marked reduction in CCR2^+^ peripheral immune cells observed following neuronal stimulation.

These results put forward neuronal activity as a critical regulator of peripheral myeloid cell identity at the onset of repair, facilitating the phenotypic conversion of innate inflammatory cells into pro-regenerative effectors, thereby widening the span of neuro-immune communication.

## Conclusion

Our work reveals neuronal activity as a potent regulator of innate immune cell phenotype during remyelination, with multifunctional effects, leading to the reduction of pro-inflammatory programs, upregulation of protective pathways, metabolic reprogramming and lipid handling. These processes converge to promote a regenerative landscape permissive to efficient myelin repair. Our study further supports the potential of neuromodulatory approaches to reshape immune responses in the injured CNS and promote regeneration and neuroprotection.

## Supporting information

Statistical Table

## Acknowledgments

We would like to thank Y. Marie and iGenseq, M. Coutelier and DAC, iQuant facility, Celis and Histomics platforms for their help in producing and analyzing the data and V. Zujovic, O. Perrot and I. Leroux for their help with RNA sequencing sample preparation. We thank Brahim Nait-Oumesmar for his critical reading of the manuscript. This research was supported by France SEP to A.D., France SEP fellowship to C.P. and INSERM and Paris Brain Institute fundings. This study was in part funded by F. Hoffmann-La Roche Ltd as part of Integrative Neuroscience Collaborations Network. The present affiliation of RR is Department of Neuromuscular Diseases, University College London, London, United Kingdom.

## Author contribution

AD and CP conceptualized the design of the study. CP, RR, MSA, PS collected data. CP, ZL, FQ and AD performed bioinformatic analysis and visualized data. VP participated in microscopy data collection and FXL in statistical analysis. AD supervised the project and wrote the manuscript with CP. BZ, BS and CL participated in writing the manuscript.

## Methods

### Animals

The care and use of mice conformed to institutional policies and guidelines (Sorbonne Université, INSERM, French and European Community Council Directive 86/609/EEC). C57bl6/J (Janvier Labs), C57BL/6Ntac-Kcnk13^em1(IMPC)H^/H (Mary Lyon Centre at MRC Harwell) and L7-ChR2-eYFP (gift from Pr C. Lena, IBENS, PSL University, Paris, France ^2^ were used for this study.

### Organotypic cultures of mouse cerebellar slices

Cerebellar slice cultures were made as previously described ^3^. Briefly, P8-10 mouse cerebella were dissected in ice cold Gey’s balanced salt solution (G9779, Sigma) complemented with 4.5 mg/ml D-Glucose (G8769-100ML, Sigma) and 1X penicillin-streptomycin (100 IU/mL, ThermoFisher Scientific), before being cut into 250μm parasagittal slices using a McIlwain tissue chopper and placed on Millicell membrane (3 to 4 slices each per animal, 0.4 μm membranes, Merck Millipore) in 50% BME (41010026, Thermo Fisher Scientific), 25% Hanks’ Balanced Salt Solution (14185-045, Thermo Fisher Scientific), 25% heat-inactivated horse serum (26050088, Thermo Fisher Scientific) medium, supplemented with GlutaMax (2 mM, 35050038, Thermo Fisher Scientific), penicillin-streptomycin (100 IU/mL, Thermo Fisher Scientific) and D-Glucose (4.5 mg/ml; Sigma). To minimize inter-animal variability inherent to *ex vivo* culture systems, slices from each animal were cultured in parallel on two separate membranes. One membrane was used for treatment, while the other served as an internal untreated control, which allowed to run more powerful statistical analysis. Cultures were maintained at 37°C under 5% CO_2_ and medium changed every two to three days. The experiments were analyzed at 10 days in vitro (10 DIV remyelination).

### Viral transduction of ex vivo cultures

To drive the expression of the DREADD receptor hM3D(Gq) in neurons, we used the adenovirus AAV8-hSyn-hM3D(Gq)-mCherry (Addgene, #50474-AAV8) as previously described ^1^. The transduction was performed immediately following slice generation, by direct deposition of the AAV solution onto the slices placed on milicell membranes (1 μl/slice at a final concentration of 1×10^11^VP/μl for AAV8-hSyn-hM3D(Gq)-mCherry).

### Optogenetic stimulation of cerebellar slice cultures

Optogenetic stimulations were performed using custom-designed set-ups allowing stimulation with a wavelength of 470nm (activation of ChR2-YFP) or 590nm (control condition) of 6-well culture plates ^1^. Each set-up contains 6 identical LEDs (SZ-05-A6 for 590nm and SZ-05-H3 for 470nm, Luxeonstar). Each LED, individually calibrated to allow a stimulation at a power of 1.5mW/mm^2^ of the slices. The pattern of stimulation was controlled with an Arduino (A000067, Arduino Mega 2560, Arduino, ref.782-A000067, Mouser Electronic). To stimulate Purkinje cells during remyelination (DIV10), organotypic cerebellar slices were subjected to either a tonic or trimodal optogenetic stimulation protocol using blue light (470 nm). For tonic stimulation, 10 ms pulses were applied at 10 Hz for either 1 hour (to study microglia–node interactions) or 6 hours (to assess microglial phenotype), followed by immediate fixation of the slices with Paraformaldehyde (PFA). The trimodal stimulation pattern consisted of 10 ms pulses at 10 Hz for 4.5 s, followed by a 100 ms continuous stimulation and a 500 ms light-off phase. This cycle was repeated continuously for 1 hour (microglia–node interaction) or 6 hours (microglial phenotype). To investigate the effect of these patterns on remyelination, the slices were stimulated for 6 hours at DIV10 and further incubated overnight before fixation the following morning.

To avoid cytotoxicity, prior to stimulation, the culture medium was replaced by a medium free of phenol-red consisting of 75% DMEM (11880028, Gibco, Thermo Fisher Scientific), 20% 1X HBSS (14185-045, Gibco, Thermo Fisher Scientific) supplemented with HCO ^-^ (0.075 g/L final; 25080060, Gibco, Thermo Fisher Scientific), 5% heat-inactivated horse serum (Thermo Fisher Scientific), HEPES Buffer (10 mM final; 15630056, Gibco, Thermo Fisher Scientific), D-Glucose (4.5 g/L final; G8769-100ML, Sigma), GlutaMax (2 mM final; 35050038, Thermo Fisher Scientific) and penicillin-streptomycin (100 IU/mL each; Thermo Fisher Scientific).

### Pharmacological treatments of cerebellar slice cultures

Demyelination was induced at 6 days in vitro (DIV6) by incubating the myelinated slices with 0.5 mg/mL L-α-Lysophosphatidylcholine (LPC; L4129, Sigma, Merck) for 16 hours in fresh culture medium.

To investigate the role of P2X7 receptor (P2X7R) signaling in microglial phenotypic modulation during remyelination under a trimodal activation paradigm, for each animal, remyelinating slices (DIV10) were exposed to 470 nm light (control condition, 6-hour stimulation), one membrane being concomitantly treated with A438079 (100 μM; MedChemExpress) to pharmacologically inhibit P2X7R during optogenetic activation phase, while the second membrane was used as control.

To activate DREADD receptors *ex vivo*, cerebellar slices were treated at 10 days in vitro (DIV10), corresponding to the remyelination phase, with 0.5 μM Clozapine-N-Oxide (CNO; #16882, Cayman Chemical; stock 5 mM in DMSO, diluted 1:1000), or with an equivalent volume of DMSO alone (0.1%, vehicle control). To study microglia–node interaction modulation by neuronal activity, the slices were treated with CNO for 1 hour and immediately fixed. To investigate microglial phenotype modulation in response to neuronal activation via hM3D(Gq), the slices were treated with CNO for 6 hours and fixed at the end of the treatment. To evaluate the impact of neuronal activation on remyelination, slices were similarly treated with CNO for 6 hours at DIV10, after which the medium was replaced with fresh medium without CNO. Slices were then maintained in culture overnight before fixation the following morning.

### Electrophysiological recordings

The electrophysiological recording of slices stimulated by optogenetics was done as previsouly described ^1^. Organotypic cerebellar slices at 9-11 DIV were transferred to a recording chamber and continuously superfused with oxygenated (95% O2 and 5% CO2) aCSF containing (in mM): 124 NaCl, 3 KCl, 1.25 NaH2PO4, 26 NaHCO3, 1.3 MgSO4, 2.5 CaCl2, and 15 glucose (pH 7.4), all from Sigma Aldrich). Purkinje cells were visualized under differential interference contrast optics using a 63X water immersion lens (N.A. 1). Loose cell-attached voltage clamp recordings of the spontaneous firing activity of Purkinje cells were performed at 30-34°C with a borosilicate glass pipette filled with aCSF. Signals were amplified with a Multiclamp 700B amplifier (Molecular devices), sampled and filtered at 10 kHz with a Digidata 1550B (Molecular Devices). Data were acquired with the pClamp software (Molecular devices). To avoid any alteration of the spontaneous firing frequency of the cell by the patch procedure, the holding membrane potential was set to the value at which zero current was injected by the amplifier. The resistance of the seal (Rseal) was controlled and calculated every minute from the current response to a voltage step (200 ms; −10 mV). Only recordings with a Rseal in the range of 10 to 100MΩ and stable during the recording procedure were included in the analysis. To test optogenetic stimulation, a LED with an excitation filter 482/35 was calibrated to stimulate the field of view at 1.5mW/mm^2^ and individual neurons were successively recorded with no stimulation and with tonic or trimodal patterns. The mean firing rate was analyzed over 110 seconds recording time window using a threshold crossing spike detection in Clampfit (Molecular devices) and calculated as the number of action potential divided by the duration of the recording.

### Fixation of cultured cerebellar slices

Cerebellar slices were fixed as described before (49), with 4% PFA (Electron Microscopy, ThermoFisher Scientific) for 5 minutes followed by 1% PFA for 25 minutes at room temperature (RT) and then washed in PBS 1X (ET330-A; Euromedex). Subsequently, the slices were incubated at −20°C for 20 minutes in absolute ethanol (Sigma Aldrich, for PLP staining) and washed in PBS 1X. The slices were next blocked for 1 hour in PBS 1X, 5% Normal Goat Serum (50-062Z; ThermoFisher Scientific), 0,3% Triton X-100 (9036-19-5, Sigma) and incubated with primary antibodies diluted in blocking solution overnight at RT. The slices were then washed in PBS 1X, incubated for 3 hours at RT in the dark with secondary antibodies diluted in blocking solution. The slices were finally washed in PBS 1X and mounted between a glass slide and a coverslip (VWR; 631-0138) with Fluoromount-G (0100-01, Southern Biotech).

### In vivo viral injection

Following an intraperitoneal injection of buprenorphine (0.1mg/mL, Buprecare, Med’Vet) 15 minutes before surgery, C57Bl6J females (8-week old) or C57BL/6Ntac-Kcnk13^em1(IMPC)H^/H (from 8 to 12-week-old) were anesthetized with isofluorane (3%/1,5% for induction/maintenance, Vetfluran). The mice were then placed onto a spinal stereotaxic frame and an incision was made to access the dorsal funiculus of the spinal cord between the thoracic and lumbar vertebrae without laminectomy. 1μL of AAVrg-hSyn-hM3D(G_q_)-mCherry (Addgene, #50474-AAVrg, 7×10^12^ vg/ml) or AAVrg-hSyn-hM4D(G_i_)-mCherry (Addgene, #50475-AAVrg, 7×10^12^ vg/ml) were then injected intraspinaly with a glass capillary (30-0041, Harvard Appartus) ^1^. Following surgery, the mice were stitched and placed into a warming chamber (Vet tech solution LTD, HE011) until recovery.

### Induction of a focal demyelination in the dorsal spinal cord

A focal demyelination of the dorsal funiculus of mouse spinal cord was induced 4 weeks after viral injection. The initial steps of the surgery were similar to the viral injection described above. An incision was made at the lumbo-thoracic level to access the dorsal funiculus of the spinal cord between two vertebrae without performing a laminectomy. An intraspinal injection of 1 μL of lysophosphatidylcholin diluted into NaCl 9‰ (LPC, 10mg/mL; Sigma-Aldrich L4129) was then performed with a glass capillary (30-0041, Harvard Apparatus). Following surgery, the mice were stitched and placed into a warming chamber (Vet tech solution LTD, HE011) until recovery.

### DREADDs receptor activation in vivo

Clozapine N-oxide (CNO; Cayman Chemical Company) was dissolved in sterile 0.9% NaCl to a final concentration of 1 mg/mL and administered intraperitoneally at a dose of 5 mg/kg.

To study microglia-node interaction, mice were euthanized 1 hour after the final CNO injection at day 9 post-LPC injection (9 dpi). To assess microglia phenotype modulation by neuronal activity, mice received two CNO injections over a 5-hour period and then euthanized.

To evaluate longer-term effects, mice received two CNO injections over a 5-hour period at 9 dpi and were euthanized at day 11 post-LPC injection (11 dpi).

### Mouse central nervous tissues fixation and collection

After receiving a Euthasol (Centravet) overdose, adult mice were perfused transcardially with 2% paraformaldehyde (PFA; Electron Microscopy Sciences). The brain and spinal cord were collected and post-fixed in PFA 2% for 30 minutes, washed in PBS 1X and incubated in PBS with 30% sucrose (S0389, Sigma Aldrich) for 2 days at 4°C for cryoprotection. The tissues were then embeded in O.C.T (Tissue-Tek, Sakura). Using a cryostat (Leica CM 1950), the brains were cut sagitally or coronally and the spinal cord cut longitudinally in 30μm and 20µm thick sections respectively. Sections were collected on Superfrost plus glass slides (48311-703, VWR). For immunohistochemistry, the slides were first placed in absolute ethanol at −20°C for 20 minutes. They were then incubated with a blocking solution containing PBS, 5% Normal Goat Serum and 0.2% Triton X-100 for at least 30 minutes at RT. Following PBS washes, the slides were incubated with the primary antibodies diluted in blocking solution overnight at RT and the next day with the secondary antibodies diluted in blocking solution for 2 hours in the dark at RT. The slides were mounted on a glass slide under a coverslip (VWR) using Fluoromount (0100-01, Southern Biotech), and left to dry at RT before being stored at 4°C.

### Antibodies

The following primary antibodies were used: mouse IgG1 anti-Pan Na_v_ (1:150-1:300, Sigma), rabbit anti-Caspr (1:300, Abcam), mouse anti-Calbindin (1:500; Sigma), rabbit anti-Calbindin (1:300; Swant), rat anti-PLP (1:10; hybridoma, kindly provided by Dr. K. Ikenaka, Okasaki, Japan), chicken anti-GFP (1:250, Millipore), chicken anti-mCherry (1:1000 to 1:2000, EnCor Biotechnology), mouse IgG2b anti-MBP (1:200; Merk; NE1019), rabbit anti-Iba1 (1:500; Wako; 019-19741), mouse IgG1 anti-Iba1 (1:200; Abcam; ab283319), mouse IgG2a anti-iNOS (1:100; BD Laboratories; 610328), goat anti-IGF1 (1:50; R&D System, AF791), rat anti-CD68 (1:200; Abcam; ab53444), mouse IgG2b anti-Spp1 (1:200; Santa Cruz Biotechnology; sc-73631), rabbit anti-P2RY12 (1:200; Alomone Labs; APR-020), rabbit anti-Cox2 (1:500; Abcam; ab15191) Secondary antibodies corresponded to goat or donkey anti-chicken, mouse IgG2a, IgG2b, IgG1, rabbit, rat coupled to Alexa Fluor 488, 594, 647 or 405 from Invitrogen (1:500 to 1:1000).

### RNAscope/Immunochemistry costainings

RNAscope was performed on remyelinating spinal cord section from mice transduced with AAVrg-hSyn-hM3D(Gq)-mCherry or mice transduced with AAVrg-hM4D(Gi)-mCherry, according to the manufacturer’s instructions. The RNAscope Multiplex Fluorescent v2 kit (ACD, #323110) was used in combination with H₂O₂ and Protease reagents (ACD, #322381), and probes targeting *Cox4i1* (#538161-C2; Bio-Techne), *Abca1* (#522251; Bio-Techne), *Il6st* (#476211-C3; Bio-Techne), *Il10ra* (#517731-C2; Bio-Techne), *Nfkb1* (431761-C3; Bio-Techne).

Spinal cords were collected at 9 days post-demyelination (DPI9), fixed in 2% paraformaldehyde (PFA), cryoprotected in 30% sucrose (S0389, Sigma-Aldrich) for 48 hours at 4 °C, and embedded in OCT compound (Tissue-Tek, Sakura), following the same protocol as described in the “Mouse central nervous tissues fixation and collection” section, then stored at –80 °C to protect RNA degradation. Before hybridization, tissues were dehydrated in a graded ethanol series: 50% ethanol for 5 min, 70% ethanol for 5 min, and 100% ethanol twice for 5 min. Samples were then treated with H₂O₂ for 10 min, followed by a 10 min incubation in target retrieval solution at 95 °C, and a final wash in 100% ethanol for 3 min. Tissue sections were air-dried and treated with Protease III for 30 min at 40 °C, then incubated with the probes for 2 h at 40 °C. Amplification steps were carried out at 40 °C as follows: Amp1 for 30 min, Amp2 for 30 min, and Amp3 for 15 min, each followed by washes. Signal detection was performed using the RNAscope Multiplex Fluorescent v2 HRP-C1, HRP-C2, or HRP-C3 detection reagents depending on the probe channel, followed by a 15 min incubation at 40 °C. Signal amplification was achieved with Opal 520 (PerkinElmer, FP1496001KT, 1:1000) diluted in RNAscope TSA Buffer (ACD, #322809) for 30 min at 40 °C. The reaction was stopped with RNAscope FL v2 Blocker for 15 min at 40 °C. Following RNAscope, immunofluorescence was performed as previously described to detect microglia, mCherry, or P2RY12 using the following primary antibodies: anti-Iba1 (1:500; Wako), anti-mCherry (1:1000; Abcam), and anti-P2RY12 (1:300; Alomone Labs).

### Confocal microscopy

Confocal imaging was performed using an inverted SP8 Leica confocal microscope equipped with 40× (NA 1.30) and 63× (NA 1.40) oil immersion objectives and controlled by LasX software (SP8, Leica, v3.5.6). Excitation was achieved using 405, 488, 565, and 647 nm laser lines.

To image organotypic cerebellar slices, regions containing Purkinje cells expressing either mCherry (chemogenetic experiments) or YFP (optogenetic experiments) were selected. Five image stacks were made by animal and condition. To assess the effect of neuronal activity on microglia–node interaction, the image stacks (160.60 μm × 160.60 μm; 1024 × 1024 pixels) were acquired using the 63× objective, with a z-step of 0.30 μm and at least 20 optical sections per stack. To evaluate the microglial phenotype in cerebellar slices, the image stacks (290.9 μm × 290.9 μm; 1024 × 1024 pixels) were acquired using the 40× objective, with a z-step of 0.35 μm and a minimum of 10 Z-sections per stack.

Imaging of the spinal cord was performed within the dorsal funiculus, centered around the peri-lesional area, as identified by DAPI and Iba1 staining. Five image stacks were acquired for each animal. To study microglia–node interaction, stacks were imaged using 63x objective (160.61 μm × 160.61 μm images; 1024 × 1024 pixels), with at least 20 optical sections per stack and a z-step of 0.30 μm. For microglial phenotype and RNAscope analyses, stacks were imaged using the 40x objective (290.9 μm × 290.9 μm images; 1024 × 1024 pixels), with at least 20 Z-sections per stack and a z-step of 0.35 μm. To visualize remyelinating areas, a mosaic image stack of the lesion (as identified by PLP and Iba1 staining) was acquired using the 40× objective (290.9 μm × 290.9 μm; 1024 × 1024 pixels). Each field comprised at least 10 Z-sections with a z-step of 0.35 μm. For quantification of nodal structure density in the mouse spinal cord, five image stacks per animal were acquired in the remyelinating lesion (160.61 μm × 160.61 μm images, 1024 × 1024 pixels), with at least 20 Z-sections and a z-step of 0.30 μm.

### Analysis

Images were anonymized using a custom script that assigned randomized filenames, concealing experimental conditions. After completing all image analyses, the files were de-anonymized to reveal their original label, and statistical analyses were subsequently performed.

### Quantification of microglia-nodes interaction ex vivo and in vivo

In cerebellar slice cultures (10 DIV) and spinal cord sections (DPI 10), microglia–node interactions were quantified using a single central optical section within each Z-stack. The remaining planes were used to confirm the nature of the structure observed (e.g., excluding axon initial segments orthogonal to the imaging plane). A contact was defined as at least one pixel positive for the microglial marker juxtaposed to or covering at least one pixel positive for the nodal marker. Quantification was performed on five fields of view per animal, and the mean percentage of nodes contacted by microglia was calculated per animal. Analyses were conducted in at least five animals per condition (from a minimum of two independent experiments for the *ex vivo* studies).

### Remyelination Index in Cerebellar Slices

To evaluate the impact of neuronal activity on remyelination, five high-resolution images per condition were analyzed per animal, each covering an entire folium (1900 × 1900 pixels, ∼550 × 550 μm). The remyelination index was computed semi-automatically using a custom ImageJ script ^4^. A region of interest including Purkinje cell axons (excluding soma and white matter tracks) was manually delineated. Binary masks were generated for the total axonal area (Calbindin+ pixels) and for the myelinated axonal area (PLP+ pixels overlapping with Calbindin+ pixels). The remyelination index was defined as the ratio of the myelinated axonal area to the total axonal area. The mean index per animal was obtained by averaging the results of the five images.

### Microglial Phenotyping (Ex Vivo and In Vivo)

To assess microglial phenotypes *ex vivo*, five fields of view (1024 × 1024 pixels, 290.9 × 290.9 μm) were analyzed per condition and per animal. The proportion of Iba1^+^ microglia expressing either IGF1 (pro-regenerative marker) or iNOS (pro-inflammatory marker) was determined by calculating the percentage of Iba1^+^ cells co-expressing each of these markers. Paired control and treated slices from each animal were analyzed. At least five animals were analyzed per condition.

For the *in vivo* experiments following LPC-induced demyelination, five images per animal were acquired at the edge of the lesion (1024 × 1024 pixels, 290 × 290 μm). The percentage of Iba1+ cells expressing IGF1, iNOS, or CD68 was quantified relative to the total Iba1^+^ population for each marker independently. Analyses were conducted in at least five animals per condition.

### Quantification of Non-Remyelinated Areas In Vivo

To assess the impact of neuronal activity on remyelination *in vivo*, we quantified the proportion of the lesion still demyelinated. For each animal, two sections were imaged using high-resolution confocal microscopy. A mosaic covering the entire lesion area was generated using the 40× objective. The total lesion area was delineated using Iba1 staining (area filled with dense ameboid Iba1+ cells). Within this area, regions lacking PLP signal were identified as non-remyelinated tissue. Area measurements were performed using ImageJ. The percentage of non-remyelinated area was calculated as the ratio of PLP⁻ area to total Iba1+ lesion area. The measurements of the two sections analyzed per animal were averaged to obtain a mean value per animal. Quantification was conducted in a blinded manner with at least five animals per condition.

### Quantification of Node density In Vivo During Remyelination

To evaluate the effect of neuronal activity on node of Ranvier formation during remyelination *in vivo*, we quantified Nav-positive clusters associated to Caspr staining (i.e., flanked by paranodal markers), which were interpreted as mature nodes. Quantification was performed on a central optical section of each Z-stack; adjacent images were examined to confirm that the structures were not heminodes or immature nodes (nodes like-clusters) extending across planes. For each field of view, the density of nodes was calculated as the number of Nav+ clusters per unit area, normalized to mm². Five images per animal were analyzed, and the mean nodal density +/- SEM per animal was calculated.

### Microglial Morphology Analysis

Microglial morphology was analyzed both *ex vivo* and *in vivo*. Z-projections were generated, and only microglia fully contained within the Z-stack were included. Images were binarized, and three to five microglia were analyzed per image (5 images per animals). Sholl analysis was performed using the Sholl plugin in ImageJ, and a mean Sholl profile was calculated per animal. A minimum of 15 microglia per animal were quantified, with at least 75 cells per condition (n ≥ 5 animals per group). For process length quantification, the binarized images were processed using the Skeletonize plugin in ImageJ, and the mean process length was calculated per microglia. A mean process length per animal was then calculated.

### RNAscope analyses

For RNAscope analysis, a central Z-plane within the stack was selected. Iba1+ microglia containing at least one nuclear punctum (based on DAPI co-staining) were counted. Two outcome metrics were calculated: the percentage of Iba1+ cells expressing the RNA of interest, and the number of puncta per positive cell. Quantification was restricted to the lesion except for *Abca1*, for which microglia were quantified outside the lesion in order to assess *Abca1* expression in P2RY12⁺ homeostatic microglia.

### Statistics

All statistical analyses and data visualizations were performed using GraphPad Prism (version 7) and R (version 2024.04.0; R Foundation for Statistical Computing, Vienna, Austria). The number of biological replicates, statistical tests used, and exact p-values are indicated in the main text, figure legends, or in the statistical summary table. Data are presented as mean ± standard error of the mean (SEM), with individual biological replicates plotted. A significance threshold of *p* < 0.05 was applied. Statistical significance is denoted as follows: *p* ≤ 0.05 (*), *p* ≤ 0.01 (**), and “ns” for non-significant differences. For experiments with sample sizes *n* ≥ 6, data distribution normality was assessed using the Shapiro–Wilk test. Parametric tests were applied when data followed a normal distribution; otherwise, non-parametric tests were used. Normality was assessed using the Shapiro–Wilk test. For experiments with *n* ≥ 6, parametric tests were applied when the data passed the normality test. For experiments with *n* < 6, non-parametric tests were used

### Bulk RNA sequencing

Eight-week-old female C57Bl/6J mice were transduced with either AAVrg-hSyn-hM3D(Gq)-mCherry, AAVrg-hSyn-hM4D(Gi)-mCherry or AAVrg-hSyn-mCherry (Addgene, #114472-AAVrg, 7×10^12^ vg/ml), followed by focal demyelination of the dorsal funiculus in the spinal cord as described above.

At 9 days post-injection (9 dpi), for each viraly-transduced group, half the mice received an intraperitoneal injection of Clozapine N-Oxide diluted in 0.9% NaCl (CNO; Cayman Chemical Company; 5 mg/kg; two injections with a 2-hour interval), while the other half was injected with 0.9% NaCl (Control). The mice were euthanized 5 hours following the first injection, and spinal cord lesioned area was collected and snap-frozen in liquid nitrogen, before being stored at −80°C. RNAs were extracted using Rneasy Mini Kit (Qiagen; 74104) according to the manufacturer instructions. RNA quality was checked using a 4200 TapeStation (G21191B) and all the samples with a RIN >7.0 were kept (mean 8.02 for mice transduced with hM3D(Gq) and 8.08 for hM4D(Gi)). RNA solutions were stored at −80°C until sequencing.

### Library preparation and sequencing of BulkRNA

Total RNA samples were used to generate mRNA libraries using the Illumina Stranded mRNA Prep kit (Illumina, CA, USA). Library preparation was performed on a LabChip GX system, including poly(A) selection, RNA fragmentation, first- and second-strand cDNA synthesis, end repair, A-tailing, adapter ligation, and PCR enrichment steps, following the manufacturer’s protocol. Library quality and concentration were assessed using the LabChip GX (PerkinElmer) for sizing and quantification, ensuring fragment size consistency and purity prior to sequencing. High-throughput transcriptome sequencing was carried out on the Illumina NovaSeq 6000 platform, using the NovaSeq 6000 S2 Reagent Kit (200 cycles), with paired-end 100 bp reads. Sequencing was performed to obtain sufficient depth for transcriptome analysis across all samples. Initial quality control of the reads was performed to evaluate sequencing quality.

### Bulk-RNA sequencing analysis

The quality of the raw data was assessed using FastQC (v0.11.5), with only high-quality paired reads retained for further analysis. The reads were then aligned to the mm10 reference genome using the Dragen pipeline, and only uniquely mapped reads were retained for downstream analysis. Gene quantification was performed using RSEM (v1.2.28) based on the RefSeq annotation, prior to normalization with the DESeq2 Bioconductor package (v3.28.0). To account for batch effect in the PCA representation, an additive model was used to include the batch variable as a covariate within the generalized linear model (GLM) framework of DESeq2. Differential expression analysis was performed using the likelihood ratio test, with a minimum log2 fold-change threshold of 0.5. We used adjusted p-value with the Benjamini–Hochberg procedure to control the false discovery rate (FDR). Functional enrichment analysis was carried out using Ingenuity Pathway Analysis (IPA, Qiagen) on the differentially expressed genes, employing an over-representation analysis approach.

### Single Cell RNA sequencing Animals

The dorsal funiculus of the spinal cord of eight-week-old female C57Bl/6J mice were transduced with either AAVrg-hSyn-hM3D(Gq)-mCherry (Addgene, #50474-AAVrg, 2.5 × 10¹³ GC/ml) or AAVrg-hSyn-hM4D(Gi)-mCherry (Addgene, #50475-AAVrg, 2.4 × 10¹³ GC/ml) to target neuronal populations projecting through this area, followed by LPC-induced focal demyelination 4 weeks later. At 9 days post-injection (9 dpi), for each virally transduced group, half the mice received an intraperitoneal injection of Clozapine N-Oxide (CNO; Cayman Chemical Company; 5 mg/kg; two injections with a 2-hour interval), while the other half was injected with 0.9% NaCl (Control). For each experiment, each experimental batch consisted of four mice, whose lesioned area of the spinal cord were pooled for tissue dissociation, with a total of four independent experiments. In total, 32 mice were transduced for each viral construct, 16 being injected with CNO and 16 with NaCl.

### Spinal cord dissociation

A 1X HBSS-based dissociation buffer was prepared using 10% HBSS without CaCl₂ and MgCl₂ (Thermo Fisher Scientific, #14185), supplemented with 1% HEPES (Thermo Fisher Scientific, #15630056), 1% sodium bicarbonate (Thermo Fisher Scientific, #25080060), 1% penicillin-streptomycin (Thermo Fisher Scientific, #15140122), and 0.5% actinomycin D (45 mM; Sigma-Aldrich, #A1410). This buffer was used throughout the protocol to rinse tissues and resuspend cells between steps and was kept on ice at all times. The enzymatic dissociation solution was prepared separately and composed of 10% 1X HBSS, 1% L-cysteine (Sigma-Aldrich, #C7880; 12.4 mg/mL), 1% Dnase I (Worthington Biochemical Corporation, #LS0021329; diluted in 0.9% NaCl), 1% papain (Worthington Biochemical Corporation), and 0.5% actinomycin D. Spinal cords from four adult mice were pooled, placed in a 40 mm Petri dish with 1 mL of enzymatic solution, finely minced with scissors and a scalpel, then transferred into a 50 mL tube using the remaining 4 mL of enzymatic solution. Tissues were incubated at 37 °C for 10 minutes in a water bath with gentle agitation every 3 minutes. Enzymatic digestion was stopped by adding 500 μL of ovomucoid protease inhibitor (Worthington, #LS003085; 7 mg/mL), followed by gentle pipetting (5–10 times) to homogenize the suspension. All subsequent steps were performed on ice or at 4 °C. The suspension was filtered through 70 µm and then 30 µm cell strainers (Miltenyi Biotec), rinsing each filter with cold 1X HBSS to limit cell loss and bringing the final volume to ∼10 mL. Cells were centrifuged at 300 g for 10 minutes at 4 °C, and the pellet was gently resuspended in 3.1 mL of cold 1X HBSS. The suspension was carefully layered over 900 μL of pre-chilled 0.9% debris removal solution (Miltenyi Biotec, #130-109-398) and overlaid with 4 mL of cold HBSS using a 1 mL tip to maintain phase separation. Samples were centrifuged at 3000 g for 10 minutes at 4 °C (maximum acceleration and braking). Upper layers were discarded, and the pellet was resuspended in 2 mL of room-temperature 1X HBSS and 4 mL of RBC lysis buffer (Roche, #11814389001). After 10 minutes of gentle manual inversion at room temperature, cells were centrifuged again at 500 g for 5 minutes at 4 °C. Finally, cells were resuspended in a small volume (<1 mL) of PBS containing 0.1% BSA (Sigma-Aldrich, #A9647) and 0.5% actinomycin D (Thermo Fisher Scientific, #20012-015) for downstream processing or FACS sorting.

### Fluorescence-activated Cell Sorting (FACS)

Cells were isolated and prepared for fluorescence-activated cell sorting (FACS) using a BD FACSAria™ Iiu cell sorter equipped with a 70 µm nozzle. Prior to sorting, cells were filtered through a 50 µm cell strainer to remove aggregates and debris. To exclude dead cells, 7-Aminoactinomycin D (7-AAD; Sigma-Aldrich, SML1633) was added to the cell suspensions at a dilution of 1:1000. Doublets were excluded based on forward scatter (FSC) and side scatter (SSC) parameters using FSC-A vs FSC-H and SSC-A vs SSC-H gating strategies. Viable, 7-AAD-negative single cells were then gated and sorted directly into PBS 1X/BSA/actinomycin D solution.

### Single-cell RNA-seq: Sample Preparation, Quality Control and Sequencing

Spinal cords were enzymatically dissociated into single-cell suspensions in PBS + 0.04% BSA. Cell concentration and viability were measured using the Bio-Rad TC20™ Automated Cell Counter with Trypan Blue 0.4% (Bio-Rad #1450013). Only samples with a viability above 60% and a sufficient number of cells were retained for library preparation. Across experiments, average viability was approximately 69%, with consistent values across NaCl and CNO-treated conditions.

Single-cell libraries were generated using the Chromium Next GEM Single Cell 3’ Reagent Kits v3.1 (10x Genomics, #20012850) and Chip G (10x Genomics, #PN-1000120) on the Chromium Controller. Reverse transcription, cDNA amplification, and library construction were performed according to the manufacturer’s protocol. The quality and quantity of amplified cDNA and final libraries were assessed using the Agilent TapeStation system, with D500 ScreenTape (Agilent #5067-5592) or High Sensitivity D1000 ScreenTape (Agilent #5067-5584), depending on the sample yield. Measured cDNA concentrations ranged from ∼2000 to over 10,000 pg/μL. Libraries were sequenced on an Illumina NovaSeq X Plus system using a 100-cycle S1 flow cell, in paired-end mode, targeting approximately 50,000 reads per cell (∼2,050 million reads per run, ∼200 Gb). Data processing was performed using Cell Ranger (v7.2.0). FASTQ files were generated with the mkfastq command, and the count function was used to generate gene-barcode matrices. Reads were aligned to the mm39 mouse reference transcriptome, excluding intronic reads.

### Single-cell RNA-seq preprocessing and analysis

Raw and filtered feature-barcode matrices were processed using a custom R pipeline. Ambient RNA contamination was removed using the decontX function (*celda* package) (v1.10.0), with raw matrices as background. Decontaminated counts were saved in 10X HDF5 format. Doublets were identified with a custom Seurat-based workflow (doublet_seurat_automata). Seurat objects were created for each sample, annotated with treatment, tissue, and doublet classification. Cells were filtered based on RNA content (25^th^–90^th^ percentiles for nFeature_RNA and nCount_RNA), <25% mitochondrial and ribosomal content, and exclusion of doublets. Cell cycle scoring and normalization were performed with Seurat’s NormalizeData and CellCycleScoring functions from the Seurat package (v4.3.0). Samples were merged, and 4,000 variable features were selected for downstream analysis. PCA was performed, and the number of relevant components was determined using the kneedle method (v0.0.1). UMAP embedding, neighbor graph construction, and clustering were performed using the top PCs. The optimal clustering resolution was selected using a custom optimal cluster function. Clusters were annotated based on canonical marker genes, and differential expression was performed using Seurat’s *FindAllMarkers*.

### Microglial subclustering and differential gene expression analysis

Microglial cells were subset from the full Seurat object) and split by sample. Each sample-specific object was independently normalized using the *NormalizeData* function and used to identify variable features via *FindVariableFeatures*. Integration features were selected using *SelectIntegrationFeatures* (nfeatures = 4000), and all sample objects were merged into a single Seurat object using the merge function. The integrated dataset was scaled using ScaleData with regression of mitochondrial gene content (percent.mt) and cell cycle scores (S.Score, G2M.Score). Principal component analysis (PCA) was performed using *RunPCA* with 100 principal components. The number of informative dimensions was selected based on the cumulative variance explained, with the minimal number of PCs required to reach 90% of total variance retained for downstream analyses. UMAP embedding (RunUMAP), k-nearest neighbor graph construction (*FindNeighbors*), and clustering (*FindClusters*) were performed using these selected dimensions. Clustering resolution was set at 0.2 based on cluster stability. Cluster-specific markers were identified using *FindAllMarkers* (minimum log₂ fold change = 0.25; minimum detection in 20% of cells), and only upregulated genes with adjusted *P* < 0.05 were retained. Differential expression between CNO- and NaCl-treated microglia was assessed using the *FindMarkers* function with the “wilcox” test. Genes were considered significant if they showed adjusted *P* < 0.05, an absolute log₂ fold change > 0.25, and were detected in ≥10% of cells in either condition. Cluster distributions across samples were also evaluated.

### Monocytes subclustering and differential gene expression analysis

Monocyte subclustering followed the same pipeline. Annotated monocyte cells were split by sample, normalized, and merged after selecting 4,000 integration features with *SelectIntegrationFeatures*. Data were scaled with regression of mitochondrial content and cell cycle scores. Dimensionality reduction (PCA), UMAP embedding, and clustering were performed using the top 100 principal components.

Cluster markers were identified using *FindAllMarkers* (log₂FC > 0.25; detection > 20%), and differential expression between treatment groups was tested with *FindMarkers* (Wilcoxon test). Genes with adjusted p-value < 0.05, |log₂FC| > 0.25, and detection in at least 20% of cells were considered significant.

### Pseudotime Analysis

To reconstruct microglial state transitions, we applied Slingshot (v2.10.0, ^5^) to the PCA-reduced space generated from Seurat (v4.3.0), specifying the cluster annotated as disease-associated microglia (DL1) as the root node. Branching was allowed by setting the multiple_trajectories parameter to TRUE.

Genes dynamically regulated along pseudotime were identified using two complementary approaches. First, generalized additive models were fitted for each gene using mgcv (v1.9-0), with expression modeled as a smooth function of pseudotime. Only cells with pseudotime ≤50 were included to focus on transition from DL to HOM clusters. We used adjusted P-values and genes with FDR < 0.01 were selected. Spearman correlations were then computed between gene expression and pseudotime rank. Genes with absolute correlation > 0.2 and high expression amplitude were retained and classified as DAM-like or HOM-like based on the evolution of expression.

To assess condition-specific effects, pseudotime distributions were compared between CNO- and NaCl-treated cells within each cluster using Wilcoxon rank-sum tests. Additionally, gene expression dynamics were modeled using condition-interaction GAMs (expr ∼ s(pseudotime, by = condition) + condition), and genes with significant interaction terms (FDR < 0.05) were considered differentially modulated by neuronal activity.

1. Ronzano R, Perrot C, Mazuir E. Neuronal activity promotes axonal node-like clustering prior to myelination and remyelination in the central nervous system. bioRxiv. 2024.
2. Chaumont J et al. Clusters of cerebellar Purkinje cells control their afferent climbing fiber discharge. Proc Natl Acad Sci U S A. 2013;110:16223–16228.
3. Thetiot M et al. Preparation and immunostaining of myelinating organotypic cerebellar slice cultures. J Vis Exp. 2019;59163.
4. Ronzano, R., et al. Microglia-neuron interaction at nodes of Ranvier depends on neuronal activity through potassium release and contributes to remyelination. Nat. Commun. 12, 5219 (2021).
5. Street K et al. Slingshot: cell lineage and pseudotime inference for single-cell transcriptomics. BMC Genomics. 2018;19:477.

