## Supplementary Figure legends for "Neuronal activity promotes repair by microglial reprogramming following demyelination"

**Figure S1 | Neuronal activity shapes microglial–node interaction and microglial phenotype depending on firing patterns**

**a,** Cerebellar slices from wild-type mice were transduced with AAV8-hSyn-hM3D(Gq)-mCherry and maintained in culture 10 days before being treated for 1 h with CNO (0.5 µM) or vehicle (DMSO). **b,** Microglial (Iba1, green) contacts at nodal structures (Nav, red) are indicated by white arrowheads. Myelin appears in gray (PLP); Percentage of nodal structures contacted by $Iba1^+$ cells. Nodal density per $mm^2$. Number of $Iba1^+$ cells per $mm^2$. **c,** Electrophysiological recordings of Purkinje cells stimulated by optogenetics in L7-ChR2-EYFP cerebellar slices (whole-cell patch-clamp). Up: tonic stimulation (10 ms ON / 90 ms OFF); Down: trimodal pattern combining 10 Hz baseline, 100ms illumination, and 500 ms silent phases. **d–e,** Distribution of instantaneous firing frequencies under tonic (**d**) and trimodal (**e**) stimulation. **f,** mean firing frequency are similar for both patterns. **g–h,** L7-ChR2-EYFP myelinated cerebellar slices were exposed to 470 nm light (Tonic or Trimodal pattern) or 590 nm light with corresponding pattern (control, Ctrl) for 1 h. **g,** Microglial (Iba1, green) contact at nodal structures (Nav, red) following Tonic stimulation are indicated by white arrowheads. Myelin appears in gray (PLP). Percentage of contacted nodes, nodal density, microglial density and process length. **h,** Microglial (Iba1, green) contacts at nodal structures (Nav, red) following Trimodal stimulation are indicated by white arrowheads. Percentage of contacted nodes, nodal density, $Iba1^+$ cell number and process length. **i–j,** L7-ChR2-EYFP demyelinated slices were stimulated with 470 nm light (trimodal pattern) for 6 h in the presence of the P2X7 receptor inhibitor A438079 (100 µM) or its vehicle (DMSO). **i,** iNOS expression (red) in microglia (Iba1, green). Percentage of $Iba1^+$ microglia expressing iNOS. **j,** IGF1 expression (red) in microglia (Iba1, green). Percentage of $Iba1^+$ microglia expressing IGF1. Data are presented as mean ± s.e.m. Paired *t*-tests were used for all comparisons; refer to Statistical Table for detailed statistics. Scale bars: **b,** 5 µm; **g–h,** 10 µm; **i–j,** 20 µm.

**Figure S2 | Neuronal activity modulates microglia–node interaction in myelinated spinal cord in vivo.**

**a–b‴,** C57BL/6J mice dorsal funiculus were transduced with a control AAVrg-hSyn-mCherry virus and mice were consequently injected with CNO or NaCl (Veh) in normal (**a**) or remyelinating (**b**) condition. **a,** Representative images of microglia (Iba1, green) contacting nodal structures (Nav, red; arrowheads indicate the contacts) and associated quantification of the percentage of nodes contacted, of nodal density (**a′**), microglial density (**a″**) and mean process length (**a‴**). **b–b‴,** Similar analysis under remyelinating conditions. **c–c‴,** In hM3D(Gq)mCherry-expressing mice, CNO treatment (1 h) significantly increased microglia–node contacts in the dorsal funiculus of the spinal cord. **c,** Representative images show microglia (Iba1, green) contacting nodes (Nav, red; arrowheads indicate contact). **c–c‴,** Quantification of the percentage of nodes contacted (**c**), of nodal density (**c′**), microglial density (**c″**) and process length (**c‴**). **d–d‴,** In hM4D(Gi)mCherry-expressing mice, neuronal inhibition reduced the proportion of microglia–node contacts in the dorsal funiculus of the spinal cord. **d,** Representative image showing decreased contacts between microglia (Iba1, green) and nodes (Nav, red). **d′–d‴,** Quantification of the percentage of nodes contacted (**d**), of nodal density (**d′**), microglial density (**d″**) and process length (**d‴**), none of which were significantly altered. Data are presented as mean ± s.e.m. Statistical analysis were made using Mann-Whitney tests, apart for Sholl analysis, performed using two-way ANOVA. Scale bars: **a–d,** 10 µm.

**Figure S3 | Transient neuronal activation induces lasting effects on microglial polarization and enhances remyelination in mouse dorsal spinal cord.**

**a,** After a 5-hour neuronal activity stimulation at the onset of repair (hM3D(Gq)mCherry-expressing mice), microglia presented an increased morphological complexity, as seen by Sholl analysis, mean process length and ramification quantification. **b,** In the hM4D(Gi)mCherry-expressing mice, neuronal inhibition led to slightly reduced microglial process length, without altering branch complexity. **c-d,** hM3D(Gq)mCherry-expressing mice received CNO or NaCl (Vehicle) at 9 days post-LPC injection (dpi) and their spinal cords were fixed 48 h later (11 dpi). **c,** Immunolabeling for myelin (PLP, grey), microglia (Iba1, green), hM3 expressing axons (mCherry, red) and cell nuclei (DAPI, blue). **c′,** Quantification of the area still deprived of myelin in the lesional area ($PLP^{-}$ fraction of the densely Iba1-labeled area), showing reduced demyelinated area in CNO-treated mice. **d–d′,** Representative images of nodal (Nav, red) and paranodal (Caspr, green) structures in the spinal remyelinating area of CNO or NaCl (Veh) treated animals. **d′,** Quantification of nodal density per $mm^2$. Data presented as mean ± s.e.m. Statistical analysis were made using Mann-Whitney tests, except for Sholl analysis, performed using two-way ANOVA. Scale bars: **c,** 50 µm; **d,** 5 µm.

**Figure S4 | THIK-1 is essential for microglia-node interaction and for activity-induced microglial modulation in vivo**

**a–a‴,** Study of WT and THIK-1 KO adult mice spinal cord in homeostatic condition. **a,** Microglia (Iba1, green) contacting nodal structures (Nav, red; arrowheads indicate node–microglia contacts). **a′,** The percentage of nodal structures contacted by microglia is significantly reduced in THIK-1 KO mice. **a″–a‴,** No differences were observed in nodal or microglial density across genotypes. **b–b″″,** Representative reconstructions of individual microglia from WT and THIK-1 KO mice. **b,** Average microglial process length was significantly reduced in THIK-1 KO mice. **b′,** Sholl analysis revealed decreased branching complexity in THIK-1 KO microglia. **b″,** Quantification of the myelinated axonal area showed no difference between genotypes. **c,** Microglia-node interaction in WT and THIK-1 KO spinal cords at the onset of repair; arrowheads indicate microglia-node contacts. **c′,** The percentage of contacted nodes is reduced in THIK-1 KO mice. **d,** Immunostainings for phagocytic (CD68, red) microglia (Iba1, green) in remyelinating spinal cords. **d′,** Thik-1 KO mice showed an increased proportion of $CD68^{+}$ $Iba1^{+}$ cells. **e–f,** Expression of iNOS (**e**) and IGF1 (**f**) (red) in microglia (Iba1, green). **e′–f′,** The proportions of $iNOS^{+}$ and $IGF1^{+}$ microglia were both significantly decreased in remyelinating spinal cord of THIK-1 KO mice. **g–j′,** THIK-1 knockout mouse spinal cords were transduced with AAVrg-hSyn-hM3D(Gq)-mCherry retrograde virus, followed by LPC-induced demyelination. The mice then received CNO or NaCl (vehicle) treatment at the onset of repair and spinal cords were collected at 9 dpi. **h,** Microglia (Iba1, green) contacting nodal structures (Nav, red) in spinal cord sections; arrowheads indicate microglia–node contacts. **h′,** Percentage of contacted nodal structures. **i–j,** Microglia (Iba1, green) expressing iNOS (**i**) or IGF1 (**j**) (red); cell nuclei are stained with DAPI (blue). **i′–j′,** Percentage of $Iba1^{+}$ cells expressing iNOS or IGF1. Data are presented as mean ± s.e.m. Mann-Whitney tests were used for all pairwise comparisons; Sholl analysis was performed using two-way ANOVA. Scale bars: **a–c,** 10 µm; **d–f,** 20 µm.

**Figure S5 | Bulk RNA-sequencing reveals a regulation of inflammatory gene expression by neuronal activity in the remyelinating mouse spinal cord.**

**a–h,** Transcriptomic profiling of spinal cord lesions collected at 9 dpi from mice transduced with AAVrg-hSyn-mCherry (**a, d**), AAVrg-hSyn-hM3D(Gq)-mCherry (**b, e, g**) or AAVrg-hSyn-hM4D(Gi)-mCherry (**c, f, h**) and treated with NaCl (Vehicle) or CNO 5 hours prior to tissue collection. **a,** Principal component analysis (PCA) of bulk RNA-seq samples from AAVrg-hSyn-mCherry-injected mice showed no segregation between NaCl-treated (orange) and CNO-treated (dark blue) groups (n = 4 per group). **b,** In contrast, PCA of hM3D(Gq)-expressing mice revealed marked separation between NaCl (light blue) and CNO-treated (light orange) samples (n = 3 per group). **c,** Similarly, hM4D(Gi)-expressing mice showed clear segregation between NaCl (yellow) and CNO (green) groups (n = 3 per group). **d,** Volcano plot showing differentially expressed genes (DEGs) in control mCherry-expressing mice (CNO vs. NaCl); only 3 genes were significantly modulated ($log_2FC > 0.5$ or $< -0.5$; FDR $< 0.05$). **e,** In hM3D(Gq)-expressing mice, neuronal activation led to the downregulation of 291 genes and the upregulation of 19 genes. **f,** In hM4D(Gi)-expressing mice, neuronal silencing induced the upregulation of 764 genes, while 120 were downregulated. **g,** Ingenuity Pathway Analysis (IPA) of DEGs in hM3D(Gq)-expressing mice indicated the downregulation of inflammatory and cytokine-related pathways, including JAK–IL-6. Key downregulated genes included *Ccl3* and *Cd9 (***i-j***)*. **h,** In contrast, IPA of DEGs in hM4D(Gi) mice revealed the activation of innate immune pathways, including the upregulation of *Ccl5*, *Tnf* and *C1qa (***k, l, m***)*. Differential expression was assessed using the Wald test (DESeq2), corrected for multiple comparisons using the Benjamini-Hochberg method (FDR < 0.05). **i-m**: Data are presented as mean ± s.e.m.

**Figure S6 | Single-cell transcriptomic profiling of mouse spinal cords following neuronal activity modulation at the onset of remyelination.**

**a,** The dorsal funiculus of C57bl6 mice spinal cord was transduced with AAVrg-hSyn-hM3D(Gq)-mCherry or AAVrg-hSyn-hM4D(Gi)-mCherry and focally demyelinated 4 weeks later by LPC injection. The mice received CNO or vehicle (NaCl) injection at 9 dpi and their spinal cords were collected 5 hours later to perform single-cell RNA sequencing. **b**, Summary table of total cell numbers, gene detection metrics, and sequencing depth. Median gene count per cell was 1,027 for hM3D(Gq) and 942 for hM4D(Gi) conditions, with over 93% of reads mapped to the genome in both datasets. **c–d**, Violin plots showing quality control metrics by cell types: total number of RNAs per cell (**c**) and the percentage of total RNA corresponding to ribosomal RNA (**d**). Microglia exhibited stable profiles across conditions. **e**, UMAP representation of all sequenced cells (n = 36,130 for hM3D(Gq); n = 56,471 for hM4D(Gi)), each color representing a cell type. **f,** Heatmap of lineage-defining marker genes across identified clusters. Expression values correspond to normalized and log-transformed counts. For each experimental group, the spinal cords of four mice were pooled per experiment, with four independent experiments (16 mice per condition).

**Figure S7 | Neuronal activity modulates inflammatory and oxidative stress-related programs in microglia at the onset of repair following demyelination.**

**a-b,** Gene Ontology enrichment (Biological Process, 2025) of significantly upregulated (**a**) and downregulated (**b**) transcripts in microglia following neuronal activity enhancement at the onset of repair (hM3D(Gq), CNO vs control). **c,** Bar plot showing representative differentially expressed microglia genes, highlighting increased expression of mitochondrial genes (*Cox4i1, Ndufa6, Atp6v1g1*) and decreased expression of inflammatory and DAM-related genes (*Spp1, Nfkb1, Lpl*) in the hM3D(Gq) condition (CNO vs control). **d,** Volcano plot of differentially expressed genes in microglia following neuronal activity inhibition at the onset of repair (hM4D(Gq), CNO vs NaCl) showing 131 genes significantly upregulated (red) and 94 downregulated genes (blue) (FDR < 0.05); ribosomal transcripts represented 71 out of 131 upregulated genes in mice expressing hM4D(Gi)-CNO vs NaCl. **e,** Bar plot showing representative differentially expressed genes, highlighting increased expression of inflammatory genes (*Cd74, C1qa and H2-D1*) and decreased expression of inflammatory and homeostatic-related genes (*Tgfrb2*) and pro-repair genes (*Zfp36*). **f,** WikiPathways enrichment analysis for upregulated transcripts after exclusion of ribosomal genes. **g,** Pathway enrichment analysis of downregulated transcripts. **h,** Microglial (Iba1 green) expression of Osteopontin (encoded by *Spp1* gene, red) at the onset of repair in the spinal cord lesion of hM4D(Gi)-expressing mice treated or not with CNO. Nuclei are in blue (DAPI). Scale bar: 20 µm. **h′,** Osteopontin expression is significantly increased in microglia in the lesion following neuronal activity inhibition at the onset of repair. Data are presented as mean ± s.e.m. Mann-Whitney test.

**Figure S8 | Neuronal activity promotes microglia phenotypic transition along a pseudotemporal trajectory toward pro-regenerative and homeostatic states**

**a,** UMAP representation of the evolution of hM3D(Gq)-transduced spinal cord microglia along pseudotime (focusing on DL1, REG and HOM microglia). Trajectory inference using Slingshot showing the progression from DAM-like microglia (DL1) to a transient pro-regenerative state (REG), leading to homeostatic (HOM) microglia. **b,** Heatmaps showing gene expression along pseudotime for representative DAM markers (top) and pro-repair and homeostatic microglial genes (bottom). **c,** Distribution of microglial cells along pseudotime in NaCl- versus CNO-treated hM3D(Gq) mice. DAM-like cells (pseudotime score < 35) were enriched in NaCl-treated mice, whereas late-stage cells (score > 35) predominated in the CNO condition. **d,** Differential gene expression analysis between CNO and NaCl conditions. A total of 69 genes were significantly upregulated and 45 downregulated (FDR < 0.05). **e–l,** Smoothed expression curves (generalized additive model with 95% confidence interval) of selected genes along pseudotime for control (Vehicle, black) and activated (CNO, red) conditions.

**Figure S9 | Neuronal activity inhibition significantly alters the microglial expression of lipid-related genes.**

**a-c** Heatmaps comparing transcript expression levels between hM4D(Gi)/CNO and hM4D(Gi)/NaCl conditions in DL1, SPL, REG and HOM microglial subclusters. **a,** Pro-inflammatory genes. **b,** Pro-repair genes. **c,** Lipid associated genes. **d,** *Abca1* transcript stainings by RNAscope (red) in P2RY12$^{+}$ microglia (green), in spinal cord sections from NaCl- or CNO-treated hM4D(Gi) mice. Cell nuclei were stained using DAPI (blue). Scale bar: 20 µm**. d′,** There is a significant decrease in the proportion of P2RY12$^{+}$ microglia expressing *Abca1* following neuronal inhibition. Data are presented as mean ± s.e.m. Mann-Whitney test.
